# IFN-γ selectively suppresses a subset of TLR4-activated genes and enhancers to potentiate M1-like macrophage polarization

**DOI:** 10.1101/437160

**Authors:** Kyuho Kang, Sung Ho Park, Keunsoo Kang, Lionel B. Ivashkiv

## Abstract

Complete polarization of macrophages towards an M1-like proinflammatory and antimicrobial state requires combined action of IFN-γ and LPS. Synergistic activation of canonical inflammatory NF-κB target genes by IFN-γ and LPS is well appreciated, but less is known about whether IFN-γ negatively regulates components of the LPS response, and how this affects polarization. A combined transcriptomic and epigenomic approach revealed that IFN-γ selectively abrogates LPS-induced feedback and select metabolic pathways by suppressing TLR4-mediated activation of gene enhancers. In contrast to superinduction of inflammatory genes *via* enhancers that harbor IRF sequences and bind STAT1, IFN-γ-mediated repression targeted enhancers with STAT sequences that bound STAT3. TLR4-activated IFN-γ-suppressed enhancers comprised two subsets distinguished by differential regulation of histone acetylation and recruitment of STAT3, CDK8 and cohesin, and were functionally inactivated by IFN-γ. These findings reveal that IFN-γ suppresses feedback inhibitory and metabolic components of the TLR response to achieve full M1 polarization, and provide insights into mechanisms by which IFN-γ selectively inhibits TLR4-induced transcription.

## Introduction

Macrophages are dynamic cells that polarize into diverse states in response to various stimuli^1–3^. M1 macrophages that are polarized by inflammatory signals, such as interferon-γ (IFN-γ) and lipopolysaccharide (LPS) play an important role in host defense against pathogens as well as the pathogenesis of chronic inflammatory diseases^4–7^. Although IFN-γ *via* Jak-STAT1 signaling pathway can induce antigen-presentation molecules and chemokines, the expression of inflammatory cytokines (NF-κB target genes), such as *TNF* and *IL6* are not activated by IFN-γ alone. In addition, LPS alone transiently activates inflammatory genes but pre-exposure to LPS rather induces tolerance and thereby resistance to subsequent Toll-like receptor (TLR) 4 stimulation. To achieve complete M1 polarization, IFN-γ primes macrophages and synergizes with LPS to activate inflammatory programs through several molecular mechanisms including chromatin remodeling and metabolic reprogramming at the level of translation^8–10^.

In the TLR4 response, autocrine IFN-β signals through STAT1-STAT2 and IRF9, which form the interferon-stimulated gene factor 3 (ISGF3) complex that binds to interferon-stimulated response elements (ISREs), to induce a feedforward loop in LPS-induced gene expression that promotes M1-like polarization^11^. On the other hand, LPS also induces feedback inhibition loops including IL-10-STAT3 anti-inflammatory pathways to prevent excessive inflammation^12^. However, the importance of over-riding feedback inhibition by IFN-γ and underlying mechanisms are not well understood. Prolonged exposure to various stimuli adjusts the responsiveness of macrophages to secondary stimulation. This feature of innate immune cells, called “innate immune memory”, plays a key role in various immune responses, for example endotoxin tolerance during sepsis, trained immunity after vaccination, and inhibition of anti-microbial responses by parasitic infections^13–16^. Priming of macrophages by IFN-γ exhibits certain similarities to training^10^. Most studies of IFN-γ priming have focused on the enhancement of secondary responses to inflammatory challenge, but IFN-γ-mediated attenuation of responses to subsequent stimulation is mostly unexplored.

Epigenomic reprogramming of macrophage-specific enhancers by a variety of micro-environmental stimuli contributes to not only the distinct phenotypes of macrophages in different tissues or diseases states, but also to innate immune memory in macrophages
^17,18^. Signal-dependent transcription factors including NF-κB, AP-1, and STATs play a critical role in dynamic changes of active enhancer landscapes in macrophages^19,20^. Our previous work demonstrated that IFN-γ priming mediates genome-wide STAT1 binding with IRF1 at *cis*-regulatory elements to increase histone acetylation to enhance the transcriptional responsiveness to subsequent LPS stimulation^8^. Unbiased transcriptome-wide analysis has revealed that environmental signals confer not only the activation of gene expression but also the repression of distinct gene sets^21–23^. Molecular mechanisms by which the key macrophage-polarization signals, such as IFN-γ or IL-4 suppress gene expression have been studied using epigenomic approaches. For example, IFN-γ can repress basal M2-like gene expression programs by two distinct mechanisms: first, IFN-γ induces the deposition of negative histone mark H3K27me3 at the promoters by EZH2 recruitment^24^, and second, IFN-γ deactivates and disassembles enhancers by suppressing binding by MAF and lineage-determining transcription factors^25^. It has also been reported that IL-4 can antagonize IFN-γ-induced transcriptional responses^26^, and directly suppress LPS-induced inflammatory responses by STAT6-dependent enhancer deactivation^27^. Despite these efforts to reveal mechanisms of transcriptional repression by IFN-γ, inhibition of LPS-inducible genes by IFN-γ, at the transcriptomic and epigenomic level has not been elucidated.

In this study, we sought to understand how IFN-γ priming selectively attenuates a component of TLR4-induced genes to fully polarize M1-like macrophages. To address this question, we performed a comprehensive transcriptomic and epigenomic analysis using primary human macrophages, which are physiologically relevant for human inflammatory disease conditions. We found that LPS-induced genes that are repressed by IFN-γ priming are subdivided into at least two subsets: those regulated by an IL-10-STAT3 negative feedback loop, and those that function in metabolic pathways and are regulated independently of IL-10. One mechanism of repression involves deactivation of LPS-induced enhancers, which similarly fall into distinct subsets. Inhibition of one subset of enhancers that harbors STAT motifs occurs via suppression of histone acetylation and recruitment of STAT3, CDK8-Mediator and cohesin. This contrasts with superactivation of TLR4-inducible genes via recruitment of STAT1 to enhancers that harbor IRF motifs. These findings provide insights into mechanisms by which IFN-γ selectively suppresses anti-inflammatory and metabolic components of the TLR response by enhancer deactivation to augment M1-like inflammatory macrophage polarization.

## Results

### IFN-γ selectively inhibits components of the LPS-induced transcriptional response

A well-established function of IFN-γ is augmentation of LPS-induced inflammatory gene expression, but little is known about the overall effects of IFN-γ on TLR4-induced transcriptional responses. To examine how IFN-γ alters LPS-induced gene expression, we performed RNA sequencing (RNA-seq) analysis of primary human macrophages cultured with or without IFN-γ for 48 hr, followed by a 3 hr challenge with LPS (**Fig. 1a**; correlation between biological replicates and principal component analysis is shown in **Supplementary Fig. 1a,b**). We focused on the 3,909 genes significantly differentially expressed (FDR < 0.01, >2-fold differences in expression) in any pairwise comparison among four conditions. *k*-means clustering classified the genes into six gene clusters that were distinctly regulated by IFN-γ and LPS (**Fig. 1b, c**). Although many of the clusters captured known and previously studied patterns of gene regulation by IFN-γ and/or LPS, there was a large subset of genes (cluster IV, n = 770) whose induction by LPS was suppressed by IFN-γ. As the negative regulation of TLR responses by IFN-γ is poorly understood, in this study we focused on class IV genes, and on enhancers that are regulated in a similar manner. As a comparison point and a control, we used class V genes (n =541) that were synergistically induced by IFN-γ and LPS and comprise canonical inflammatory genes such as *IL6*, *IL23A* and *CXCL9*.

**Fig. 1.**
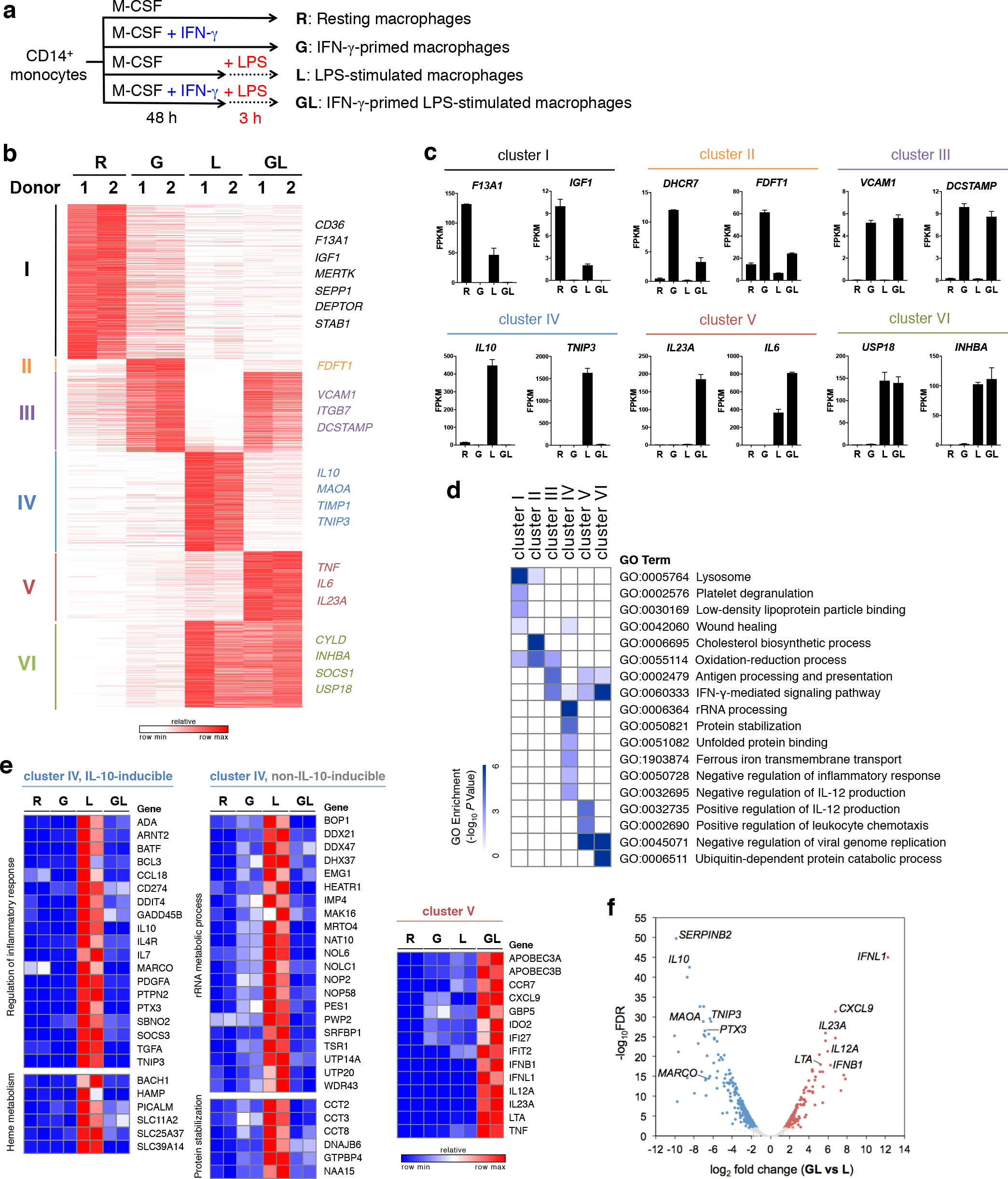
IFN-γ-mediated reprogramming of TLR4-induced transcriptome in human macrophages. (**a**) Experimental design: human CD14^+^ monocyte-derived macrophages underwent one of four different combinations of priming for 48 hr with IFN-γ before stimulation with LPS (or not) for 3 hr: without priming or stimulation (R); primed with IFN-γ without LPS stimulation (G); LPS stimulation only with no priming (L); or primed with IFN-γ and stimulated with LPS (GL). (**b**) K-means (K = 6) clustering of 3,909 differentially expressγed genes in any pairwise comparison among four conditions. Clusters are indicated on the left. (**c**) Examples of expression of select genes from the six clusters identified in (**a**). (**d**) Heat map showing the *P* value significance of GO term enrichment for genes in each cluster. (**e**) Heatmap showing the relative expression of representative cluster IV genes that are inducible by IL-10 (left), cluster IV genes that are not inducible by IL-10 (middle), and cluster V genes. Distinct biological functions are indicated on the left. IL-10-inducible genes were obtained from GSE43700. (**f**) Volcano plot of transcriptomic changes between LPS (L) and IFN-γ-primed LPS-stimulated (GL) macrophages; colored dots correspond to genes with significant (FDR < 0.01) and greater than two-fold expression changes. Data from two independent experiments with different donors are depicted.

Gene ontology (GO) analysis revealed that each cluster was enriched in genes associated with distinct biological functions (**Fig. 1d**). Cluster I, which contains genes basally expressed in human macrophages and repressed by IFN-γ and LPS, was enriched for genes involved in wound healing and related reparative processes; this extends our previous work showing that IFN-γ suppresses basal expression of genes that are inducible by glucocorticoids and IL-4^25^. In contrast, LPS-inducible genes repressed by IFN-γ (cluster IV) were associated with negative regulation of inflammatory responses, metabolism, and iron transport (**Fig. 1d**). Closer examination of these genes and comparison to public databases (GSE43700^28^) revealed that cluster IV contains *IL10* and that approximately 30% (239/770) of genes in cluster IV correspond to IL-10-inducible genes (**Fig. 1e**, left, representative genes are shown, and **Supplementary Fig. 1c**). These results suggest that IFN-γ broadly interrupts the IL-10-mediated LPS-induced negative feedback loop that negatively regulates inflammation, at least in part by suppressing *IL10* induction. GO and pathway analysis revealed that the IL-10-inucible genes in cluster IV were associated with anti-inflammatory and heme metabolism pathways, whereas the non-IL-10-inducible genes showed enrichment in distinct pathways related to lipid, purine, tryptophan, and iron metabolism; rRNA processing; and protein stabilization and unfolded protein binding (**Fig. 1e**, middle, representative genes are shown, **Supplementary Fig. 1d-f**). The two distinct gene subsets in cluster IV could be partially distinguished by induction mediated by Jak-STAT signaling versus mTORC1 signaling and Myc binding at promoters, which is in accord with a previous report^9^ (**Supplementary Fig. 1f,g**). Cluster IV also includes AP-1-dependent genes such as MMPs, which is consistent with previous reports that IFN-γ suppresses AP-1 pathways^6,29^. In contrast to class IV, class V, which contains genes synergistically induced by IFN-γ and LPS, was enriched in inflammatory cytokine and chemokine genes (**Fig. 1c-e**). The differential regulation of genes in clusters IV and V by IFN-γ is highlighted in the volcano plot depicted in **Fig. 1f**. These results identify two groups of LPS-inducible genes that are regulated in opposing directions by IFN-γ and have distinct functions, and identify negative regulation of LPS-induced metabolic genes as a new IFN-γ function.

### IFN-γ selectively inhibits a subset of LPS-activated enhancers

We wished to understand mechanisms underlying suppression of LPS-inducible genes by IFN-γ and tested the hypothesis that IFN-γ inhibits a subset of LPS-activated enhancers using an epigenomic approach. We defined active enhancers as regions of “open” chromatin (detected by ATAC-seq) that were located >1 kb away from the TSS, bound lineage-determining transcription factors (TFs) PU.1 and/or C/EBP, and exhibited histone 3 lysine 27 acetylation (H3K27-Ac) in at least one of the four conditions (FDR < 0.05, >2-fold changes in any pairwise comparison; n = 20,782 enhancers, **Supplementary Fig. 2b**). The enhancer calls were validated relative to DNase-seq and histone 3 monomethylated at lysine 4 (H3K4me1) data for CD14-positive monocytes from ENCODE project (**Supplementary Fig. 2c**). Similar to the transcriptomic data, enhancers differentially regulated at the H3K27-Ac level sorted into six major clusters (**Supplementary Fig. 2a, d** and **Fig. 2a-c**). The patterns of H3K27-Ac regulation in enhancer clusters generally resembled patterns of gene expression in gene clusters. Notably, we detected enhancer clusters whose activation (defined by increased H3K27-Ac) by LPS was either blocked (cluster e4, similar pattern to gene cluster IV) or superinduced (cluster e5, similar pattern to gene cluster V) by IFN-γ (**Fig. 2a,b**, representative gene tracks shown in **Fig. 2c**). Thus, IFN-γ also differentially regulates LPS-mediated enhancer activation. We next tested the relationship between IFN-γ-mediated changes in LPS-induced enhancer and gene activity. Notably, genes associated with enhancer cluster e4 closely correlated with genes in cluster IV, and genes associated with e5 enhancers correlated with cluster V genes (**Fig. 2d**). These enhancer-associated genes exhibited the expected pattern of gene expression, namely attenuation (e4-associated genes) or superinduction (e5-associated genes) of the LPS response by IFN-γ (**Fig. 2e,f**). Overall, these findings suggest that IFN-γ selectively modulates LPS-induced enhancer activation to attenuate or boost expression of different subsets of LPS-inducible genes.

**Fig. 2.**
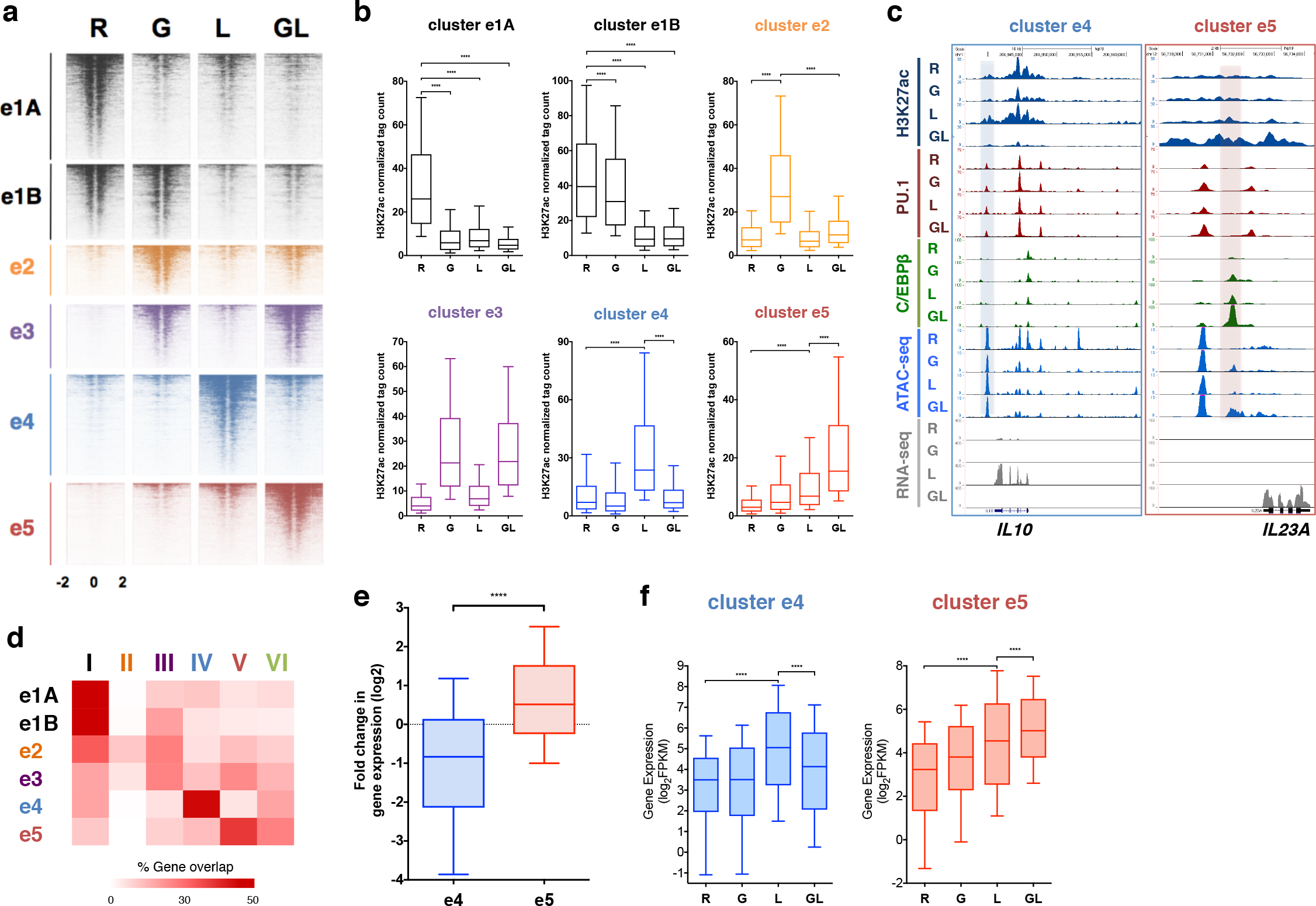
Selective regulation of LPS-activated enhancer landscapes by IFN-γ. (**a**) K-means clustering (K = 5) of enhancers as shown in Supplementary Fig. 2a further filtered (FDR < 0.05, >2-fold changes) and subdivided into six enhancer clusters. Heatmaps showing H3K27ac ChIP-seq signals at each enhancer cluster. (**b**) The boxplots indicate normalized tag counts at each enhancer cluster. ****p < 0.0001, paired-samples Wilcoxon signed-rank test. (**c**) Representative UCSC Genome Browser tracks displaying normalized tag-density profiles at enhancers of *IL10* and *IL23A* in four conditions. Shaded boxes enclose a cluster e4 enhancer (left) and cluster e5 enhancer (right). (**d**) Heatmap presentation of the percentage of genes in each cluster (**Fig. 1a**) that overlap with genes associated with the differentially regulated enhancer clusters identified in panel **a**. (**e**) Boxplots showing the change in gene expression between LPS and IFN-γ-primed and LPS-stimulated macrophages for the differentially expressed genes nearest (within 100 kb) to cluster e4 or e5 enhancers. ****p < 0.0001 by Welch’s t test. (**f**) Boxplots showing gene expression in the four indicated conditions for e4-or e5-associated genes. ****p < 0.0001 by paired-samples Wilcoxon signed-rank test. Boxes encompass the 25th to 75th percentile changes. Whiskers extend to the 10th and 90th percentiles. The central horizontal bar indicates the median. Data is representative of two independent experiments.

### TF expression and binding motif enrichment in distinct enhancer clusters

We reasoned that patterns of TF expression under the four experimental conditions, combined with TF binding motif enrichment in different enhancer clusters, would provide insight into differential regulation of enhancer activity by LPS- and IFN-γ-regulated TFs. IFN-γ and LPS altered the expression of 342 TFs which were partitioned into the six gene clusters defined in **Fig. 1b** based on pattern of expression (**Fig. 3a, b**). Cluster IV TFs (IFN-γ attenuates LPS-induced expression) were distinguished from cluster V TFs (IFN-γ synergizes with LPS) by expression of *STAT3* and AP-1 family members *BATF* and *FOSL2*. A parallel analysis of TF binding motifs enriched under the enhancer peaks defined in **Fig. 2b** revealed that enhancer clusters e4 and e5 were distinguished by enrichment of AP-1 and STAT motifs in e4 enhancers and IRF motifs in e5 enhancers (**Fig. 3c**, known motif analysis, **Fig. 3d-e**, *de novo* motif analysis, and **Supplementary Fig. 3a,b**). The enrichment of AP-1 motifs in e4 enhancers is consistent with inhibition of AP-1 signaling by IFN-γ^6,30^. Given that IFN-γ strongly induces and activates STAT1 (**Fig. 3b**), which has a predominantly pro-inflammatory function, the absence of STAT motifs in e5 enhancers may appear puzzling. However, this can be explained by previous work from our laboratory and other groups showing that treatment with IFN-γ for the longer times used in this study results in predominant binding of STAT1 to IRF sites^8,20,26,31^, presumably in a complex with IRF proteins; STAT1 binding measured by ChIP-seq experiments is analyzed below. Known motif analysis (**Fig. 3c**) suggested binding of STAT3 to e4 enhancers under LPS-stimulated conditions, which is consistent with the well-established LPS-induced IL-10-STAT3 autocrine loop, and was further supported by Factorbook ChIP-seq data from ENCODE project showing co-occupancy of STAT3 with AP-1 at representative cluster e4 enhancers (**Fig. 3f**). Furthermore, IFN-γ suppressed LPS-induced STAT3 expression (**Fig. 3b**), and ChIP-qPCR showed that IFN-γ blocked LPS-induced recruitment of STAT3 and the AP-1 protein c-Jun to e4 enhancers at the *IL10* locus (**Fig. 3g** and **h**). Overall, these data suggested a role for STAT3 in the regulation of e4 enhancers, and motivated further investigation into how the genomic profile of STAT3 binding is affected by LPS and IFN-γ.

**Fig. 3.**
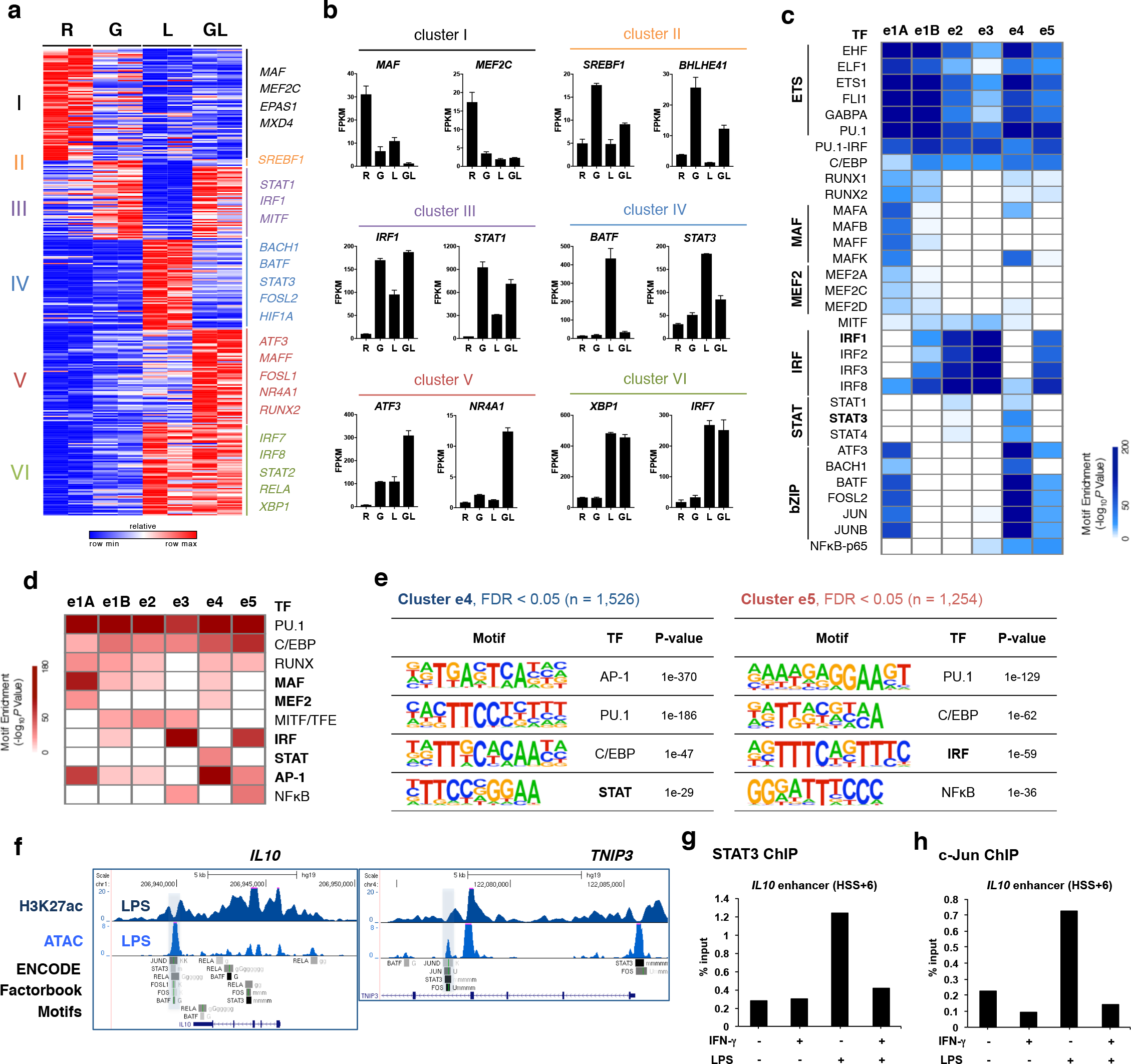
TF expression and binding motif enrichment in distinct enhancer subsets. (**a**) Heatmap of gene expression of 91 transcription factors in the clusters defined in **Fig. 1a**. (**b**) Examples of TF gene expression from the six identified clusters identified in (**a**). (**c**) Heat map showing the *P* value significance of known motif enrichment in each cluster (defined as in **Fig. 2a**) and grouped according to TF families (left). (**d**) Heat map showing the *P* value significance of *de novo* motif enrichment in six enhancer clusters. (**e**) The most significantly enriched transcription factor (TF) motifs identified by *de novo* motif analysis using HOMER in cluster e4 (left) and cluster e5 (right) enhancers. (**f**) UCSC Genome Browser tracks showing normalized tag density of H3K27ac ChIP-seq and ATAC-seq in LPS-stimulated macrophages. Cumulative TF binding at each e4 enhancer from ENCODE project (Factorbook) is shown below the gene tracks. Boxes enclose representative e4 enhancers at *IL10* (left) and *TNIP3* (right) locus. (**g**) ChIP-qPCR of STAT3 at the e4 enhancer (HSS+6) of *IL10*. (**h**) ChIP-qPCR of c-Jun at the e4 enhancer (HSS+6) of *IL10*. Data depict experiments with two different donors (**a**) or are representative of two (**b-f**) or three (**g, h**) independent experiments.

### e4 enhancers bind STAT3 which is suppressed by IFN-γ

We next performed STAT3 ChIP-seq experiments and analyzed binding of STAT3 and STAT1 (GSE43036^8^) to enhancers in clusters e1-e5 after stimulation of human macrophages with LPS, IFN-γ, or both, under the same conditions as used in **Figures 1–3**. Interestingly, induction of STAT3 occupancy showed a very restricted pattern, with a substantial increase only in cluster e4 enhancers in cells stimulated by LPS; this increase in binding was strongly blocked by IFN-γ (**Fig. 4a-c**; representative gene tracks at *IL10*, *TNIP3*, and *IL4R* loci are shown in **Fig. 4d**). The much weaker recruitment of STAT3 to e5 enhancers was not affected by IFN-γ, which is consistent with the different regulation of e4 and e5 enhancers. In contrast to STAT3, STAT1 occupancy was increased in most enhancer clusters in cells treated with IFN-γ, and was also induced by LPS in clusters e4 and e5 (**Fig. 4e-g**). As expected^8,10^, the pattern of STAT1 binding was similar to IRF1 binding (**Supplementary Fig. 4b,c**). The significance of STAT1 binding to e4 enhancers is not clear, but the increased binding of STAT1 to e5 enhancers in macrophages treated with IFN-γ + LPS is consistent with our previously proposed model of activation of ‘synergy genes’ by concomitant binding of STAT1 to several enhancers at these gene loci^8^. Overall, the data support a role for IFN-γ-mediated suppression of STAT3 binding at e4 enhancers in the downregulation of expression of associated genes.

**Fig. 4.**
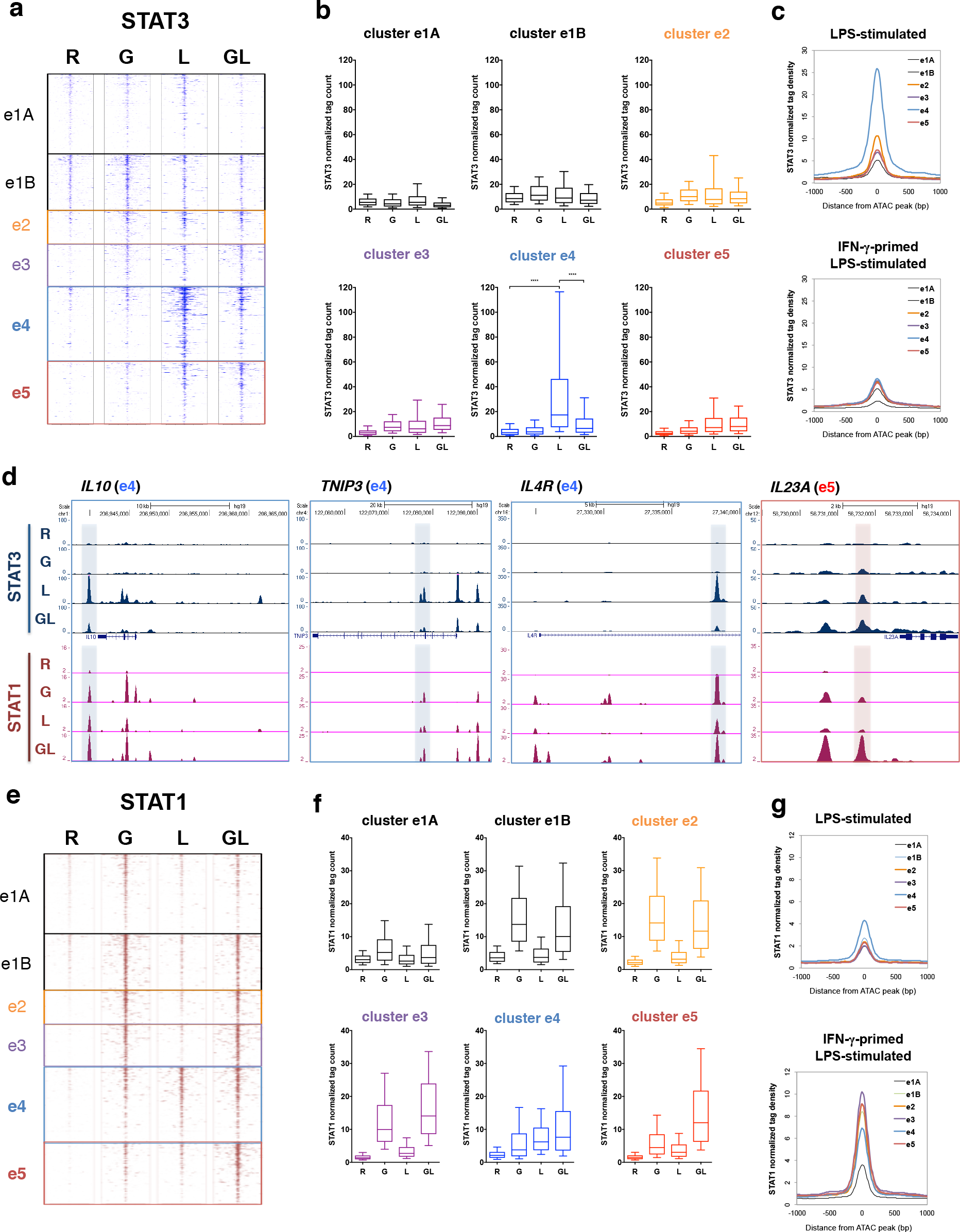
Differential occupancy of STAT3 and STAT1 at e4 and e5 enhancers. (**a**) Heatmaps showing STAT3 ChIP-seq signals at each enhancer cluster defined in **Fig. 2a**. (**b**) Boxplots depicting normalized tag counts at each enhancer cluster. ****p < 0.0001, paired-samples Wilcoxon signed-rank test. (**c**) Distribution of the average signal of STAT3 ChIP-seq at each enhancer cluster in LPS-stimulated (top) and IFN-γ-primed LPS-stimulated macrophages (bottom). (**d**) Representative UCSC Genome Browser tγracks displaying normalized tag-density profiles at enhancers of *IL10, TNIP3, IL4R,* and *IL23A* in the four indicated conditions. Boxes enclose cluster e4 enhancer (blue) and cluster e5 enhancer (red). (**e**) Heatmaps showing STAT1 ChIP-seq signals at each enhancer cluster. (**f**) The boxplots indicate normalized tag counts at each enhancer cluster. (**g**) Distribution of the average signal of STAT1 ChIP-seq at each enhancer cluster in LPS-stimulated (top) and IFN-γ-primed LPS-stimulated macrophages (bottom). Data are representative of two independent expγeriments each of which included at least two independent donors (**a-d**) or are from GSE43036 (**e-g**).

### IFN-γ suppresses coactivator and CDK8 recruitment to e4 enhancers

STATs activate gene expression in part by recruiting transcriptional coactivators such as the histone acetyltransferase p300 and Mediator complexes, a subset of which contain the serine kinase CDK8^32–34^. CDK8 in turn potentiates the transcriptional activity of STAT1 and STAT3 by phosphorylating a serine residue in their transactivation domains. We wished to test the idea that IFN-γ suppresses coactivator recruitment to e4 enhancers (LPS-activated enhancers suppressed by IFN-γ). We first examined coactivator and CDK8 recruitment at the prototypical e4 enhancer associated with the class IV gene *IL10*; as a contrasting control, we used e5 enhancers associated with prototypical class V ‘synergy genes’ genes *IL6* and *IL23A*. IFN-γ strongly suppressed LPS-induced recruitment of p300, the MED1 component of Mediator, and CDK8 to *IL10* locus e4 enhancers but not to *IL23A* and *IL6* e5 enhancers (**Fig. 5a-c** and **Supplementary Fig. 5a,b**). We followed up these results using ChIP-seq to obtain the genomic profile of CDK8 binding. Strikingly, CDK8 recruitment was most prominent at e4 and e5 enhancers and paralleled their activity (**Fig. 5d-f** and **Supplementary Fig. 5c,d**). Namely, LPS-inducible recruitment of CDK8 to e4 enhancers was suppressed by IFN-γ, and to e5 enhancers was increased by IFN-γ. Taken together, the data suggest that IFN-γ suppresses recruitment of STAT3 and coactivators to enhancers concomitant with suppressing expression of associated genes.

**Fig. 5.**
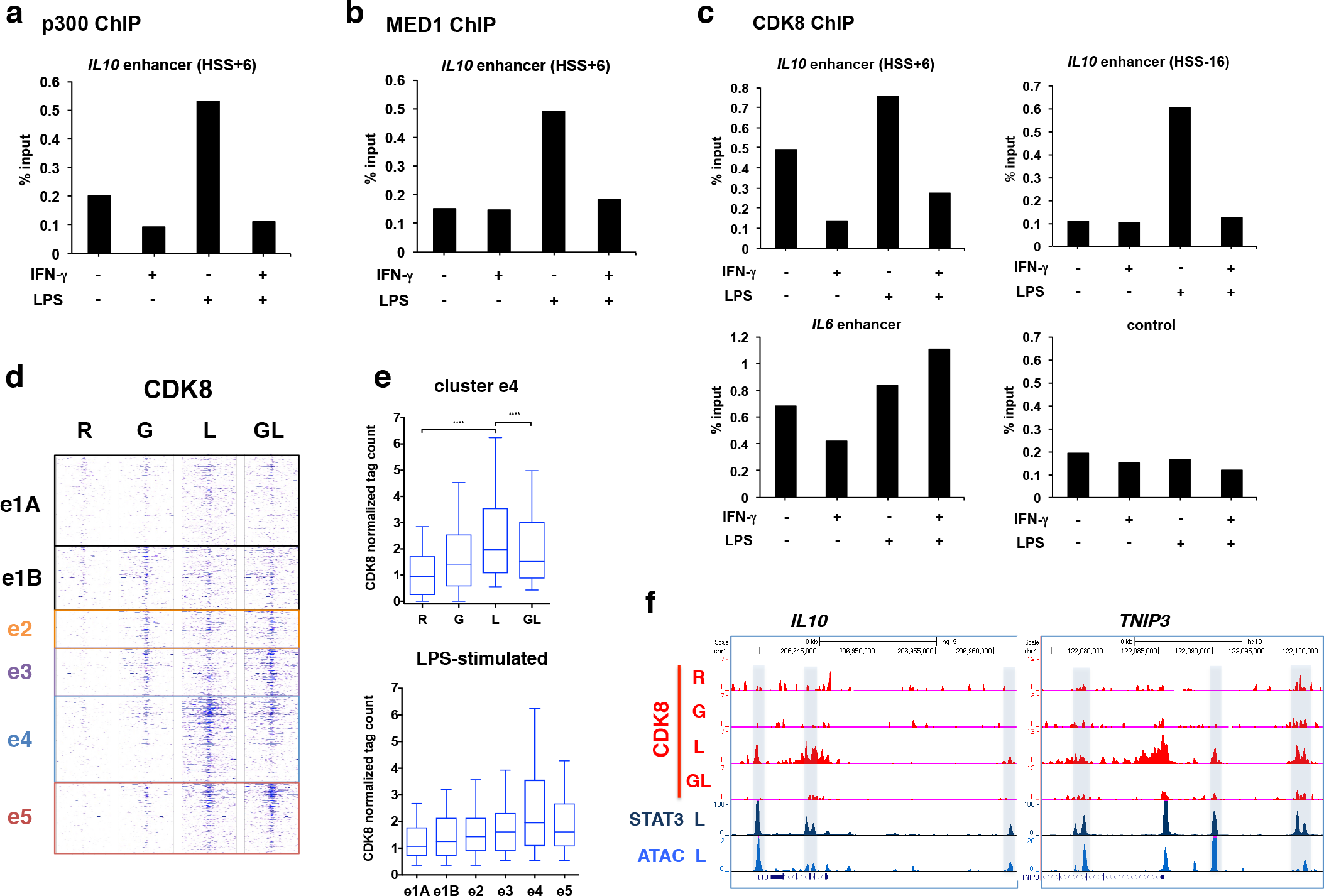
IFN-γ suppresses coactivator and CDK8 recruitment to e4 enhancers. (**a**) ChIP-qPCR analysis of p300 occupancy at the e4 enhancer (HSS+6) of *IL10*. (**b**) ChIP-qPCR analysis of MED1 occupancy at the e4 enhancer (HSS+6) of *IL10*. (**c**) ChIP-qPCR analysis of CDK8 occupancy at e4 enhancers (HSS+6 and HSS−16) of *IL10* and e5 enhancer of *IL6*. (**d**) Heatmap showing CDK8 ChIP-seq signals at each enhancer cluster defined in **Fig. 2a**. (**e**) The boxplot (top) indicates normalized tag counts at e4 enhancer in the 4 indicated conditions. ****p < 0.0001, paired-samples Wilcoxon signed-rank test. The boxplot (bottom) indicates normalized tag counts at each enhancer cluster in LPS-stimulated macrophages. (**f**) Representative UCSC Genome Browser tracks displaying normalized tag-density profiles at e4 enhancers of *IL10* and *TNIP3* in the four indicated conditions (CDK8) and the LPS-stimulated condition (STAT3 ChIP-seq and ATAC-seq). Data are representative of three independent experiments (**a-c**), or depict one ChIP experiment using pooled samples from independent experiments using four different donors (**d, e**).

### e4 enhancers partition into two subsets based on coordinate STAT3 and CDK8 binding

As cluster IV genes partitioned into IL-10-dependent and independent genes, we wondered whether e4 enhancers can be similarly subdivided. Closer examination of STAT3 binding to e4 enhancers, when rank-ordered according to tag counts, revealed that e4 enhancers partitioned into two groups, characterized by either high (52%, n =787) or low (48%, n = 739) if not absent STAT3 binding (**Fig. 6a**). We wondered whether STAT3^hi^ peaks corresponded to enhancers that are activated by autocrine IL-10, and began to address this question by performing STAT3 ChIP-seq using macrophages stimulated with recombinant IL-10. Interestingly, STAT3^hi^ e4 enhancers (in cells treated with LPS) corresponded closely to enhancers that bound STAT3 after stimulation with IL-10 (**Fig. 6b**). To determine which additional features distinguished STAT3^hi^ from STAT3^lo^ e4 enhancers, we performed motif analysis, which revealed selective enrichment of the STAT motif only at STAT3^hi^ e4 enhancers (**Fig. 6c**). This result coincides with selective enrichment of the STAT motif in promoters of IL-10-inducible cluster IV genes (Supplementary **Fig. 1g**), which showed higher STAT3 occupancy than promoters of non-IL-10-inducible cluster IV genes (**Supplementary Fig. 6a**). Furthermore, STAT3^hi^ e4 enhancers showed higher amounts of CDK8 binding compared to STAT3^lo^ e4 enhancers (**Fig. 6d**). In contrast, CDK8 occupancy was significantly higher at STAT1^hi^ e5 enhancers in IFN-γ-primed LPS-stimulated macrophages (**Supplementary Fig. 6b,c**). Thus, consistent with the literature, CDK8 occupancy paralleled STAT recruitment during macrophage activation.

**Fig. 6.**
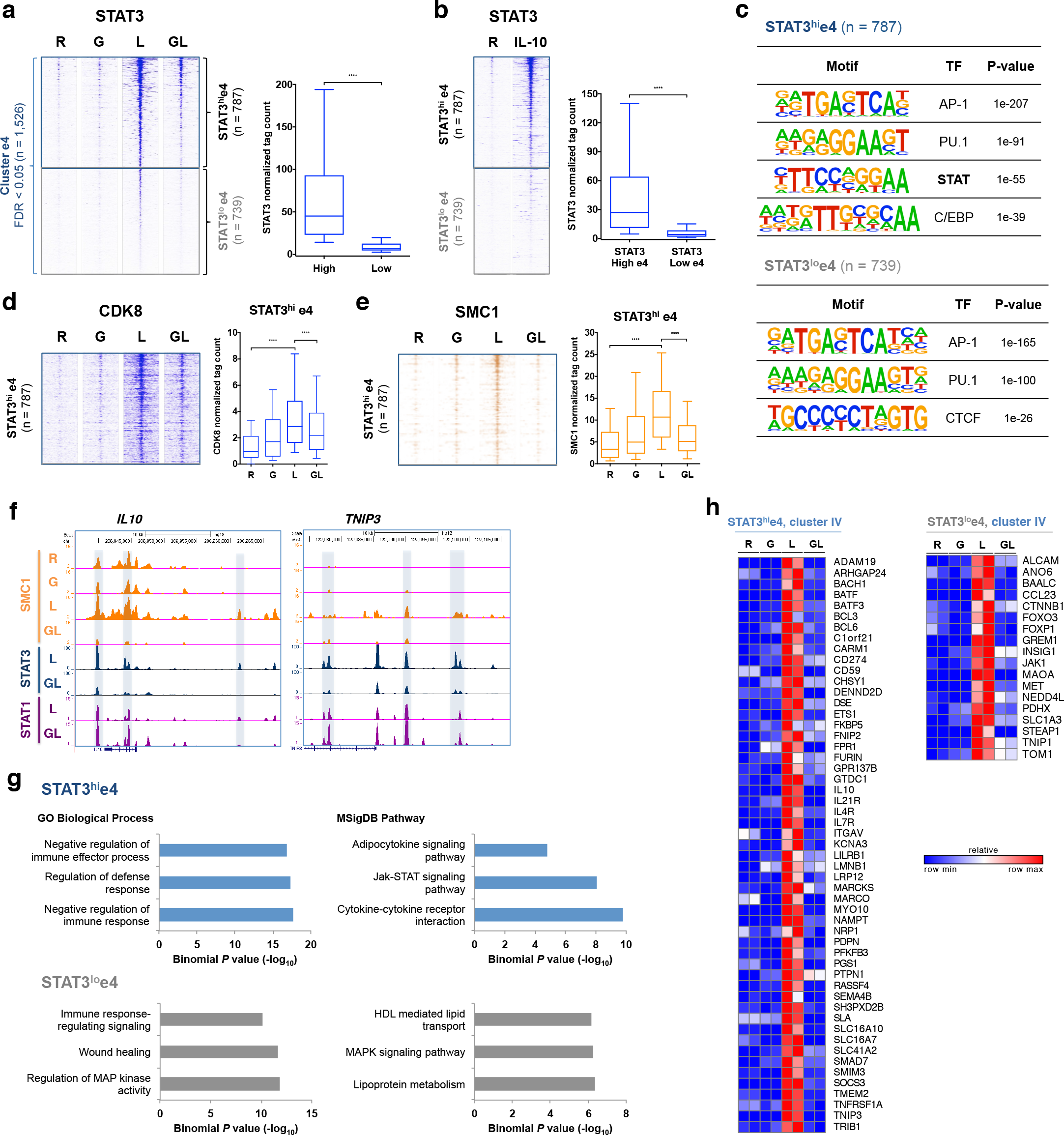
The strength of STAT3 binding divides e4 enhancers into two subgroups. (**a**) Heatmaps of STAT3 ChIP-seq signals at cluster e4 enhancers in the four indicated conditions. Enhancers were separated into two subsets: STAT3^hi^e4 (n = 787) and STAT3^lo^e4 (n = 739) based upon a cutoff of log_2_ normalized tag counts = 3). The boxplot (right) depicts normalized tag counts at STAT3^hi^e4 and STAT3^lo^e4 enhancers. (**b**) Heatmaps of STAT3 ChIP-seq signals at two subsets of e4 enhancers (defined in **a**) in resting and IL-10-stimulated macrophages. The boxplot (right) indicates normalized tag counts at STAT3^hi^e4 and STAT3^lo^e4 enhancers. (**c**) The most significantly enriched transcription factor (TF) motifs identified by *de novo* motif analysis using HOMER at STAT3^hi^e4 (top) and STAT3^lo^e4 (bottom) enhancers. (**d**) Heatmaps of CDK8 ChIP-seq signals at STAT3^hi^e4 enhancers in the four indicated conditions. The boxplot (right) indicates normalized tag counts at STAT3^hi^e4 enhancers. (**e**) Heatmaps of SMC1 ChIP-seq signals at STAT3^hi^e4 enhancers in the four indicated conditions. The boxplot (right) indicates normalized tag counts at STAT3^hi^e4 enhancers in four conditions. ****p < 0.0001, paired-samples Wilcoxon signed-rank test. (**f**) Representative UCSC Genome Browser tracks displaying normalized tag-density profiles at e4 enhancers of *IL10* and *TNIP3* in the indicated conditions. (**g**) Enriched Gene Ontology (GO) and MSigDB pathway categories of genes assigned to STAT3^hi^e4 enhancers (upper panel) or STAT3^lo^e4 enhancers (lower panel). (**h**) Heatmaps of IL-10-inducible cluster IV genes that correspond to STAT3^hi^e4-associated genes (left panel) or non-IL-10-inducible cluster IV genes that correspond to STAT3^lo^e4-associated genes (right panel). Data are representative of two independent experiments each of which included at least two independent donors (**a, c, g, h**) or depict one ChIP experiment using pooled samples from independent experiments using two (**b**) or four different donors (**d, e**).

CDK8-Mediator complexes and cohesin co-occupy regulatory elements involved in enhancer-promoter looping during gene activation^35–37^. We thus wished to investigate the relationship between CDK8 and cohesin occupancy in STAT3^hi^ and STAT3^lo^ e4 enhancers. Similar to CDK8 binding, SMC1 occupancy was higher at STAT3^hi^ e4 enhancers, and this binding was induced by LPS and suppressed by IFN-γ (**Fig. 6e,f**; representative gene tracks at *IL10* and *TNIP3*). We next tested whether the two distinct groups of e4 enhancers (STAT3^hi^ CDK8^hi^ SMC1^hi^ *versus* STAT3^lo^ CDK8^lo^ SMC1^lo^) are associated with the different subsets of cluster IV genes, as defined above in **Fig. 1**. Genomic Regions Enrichment of Annotations Tool (GREAT) analysis of genes associated with STAT3^hi^ e4 enhancers revealed enrichment for IL-10-related GO terms and pathways (**Fig. 6g**, upper panels and 6h), similar to IL-10-inducible cluster IV genes (**Fig. 1e** and **Supplementary Fig. 1e, f**). In contrast, genes associated with STAT3^lo^ e4 enhancers showed distinct functional enrichment for lipid metabolism and MAPK signaling pathways (**Fig. 6g**, lower panels), which partially resembles pathways associated with non-IL-10-inducible cluster IV genes (**Supplementary Fig. 1d-f**) and is consistent with enrichment of AP-1 motifs in promoters and enhancers of these genes (**Supplementary Figs. 1f, g** and **6c**). Indeed, genes associated with STAT3^hi^ e4 enhancers were highly represented in the IL-10-inducible cluster IV gene set (53 out of 101; **Fig. 6h**, left), whereas genes associated with STAT3^lo^ e4 enhancers were included in the non-IL-10-inducible cluster IV gene set (18 out of 36; **Fig. 6h**, right). In addition, genes associated with STAT3^hi^ e4 enhancers were more highly induced by IL-10 in monocytes (**Supplementary Fig. 6d**). Overall, these data suggest that, similar to cluster IV genes, e4 enhancers partition into two groups: those activated by LPS-induced autocrine IL-10 that bind STAT3 as well as CDK8-Mediator and cohesin, and those possibly regulated by AP-1 or other as yet unknown IL-10-independent mechanisms.

### IFN-γ functionally suppresses LPS-inducible STAT3-binding enhancers

Enhancer activity is associated with recruitment of RNA polymerase II (Pol II) and transcription of enhancer RNAs (eRNAs)^38–41^. To corroborate the notion that IFN-γ inactivates LPS-induced enhancers, as suggested by the analysis of histone acetylation and CDK8-cohesin recruitment presented above, we more directly measured enhancer activity using ChIP-seq to measure Pol II recruitment. IFN-γ attenuated LPS-induced Pol II recruitment to STAT3^hi^ e4 enhancers (**Fig. 7a, b**; representative gene tracks at *IL10* locus; HSS+6 and HSS−16 mark enhancers). In contrast, IFN-γ increased Pol II recruitment to STAT1^hi^ e5 enhancers (**Supplementary Fig. 7a, b**; representative gene tracks at *IL23A* locus). eRNA could not be reliably measured genome-wide by RNA-seq, but IFN-γ-mediated suppression was clearly seen at enhancers of select genes such as *IL10* (**Fig. 7b**, gene tracks 5-8) and was confirmed by qPCR (**Fig. 7c** and **Supplementary Fig. 7c, d**). Locked nucleic acid-mediated knockdown of *IL10* eRNA at enhancers located 16 kb upstream (HSS−16) or 6 kb downstream (HSS+6) of the *IL10* TSS suppressed *IL10* mRNA expression, supporting a functional role for eRNA and for IFN-γ-regulated enhancer activity.

**Fig. 7.**
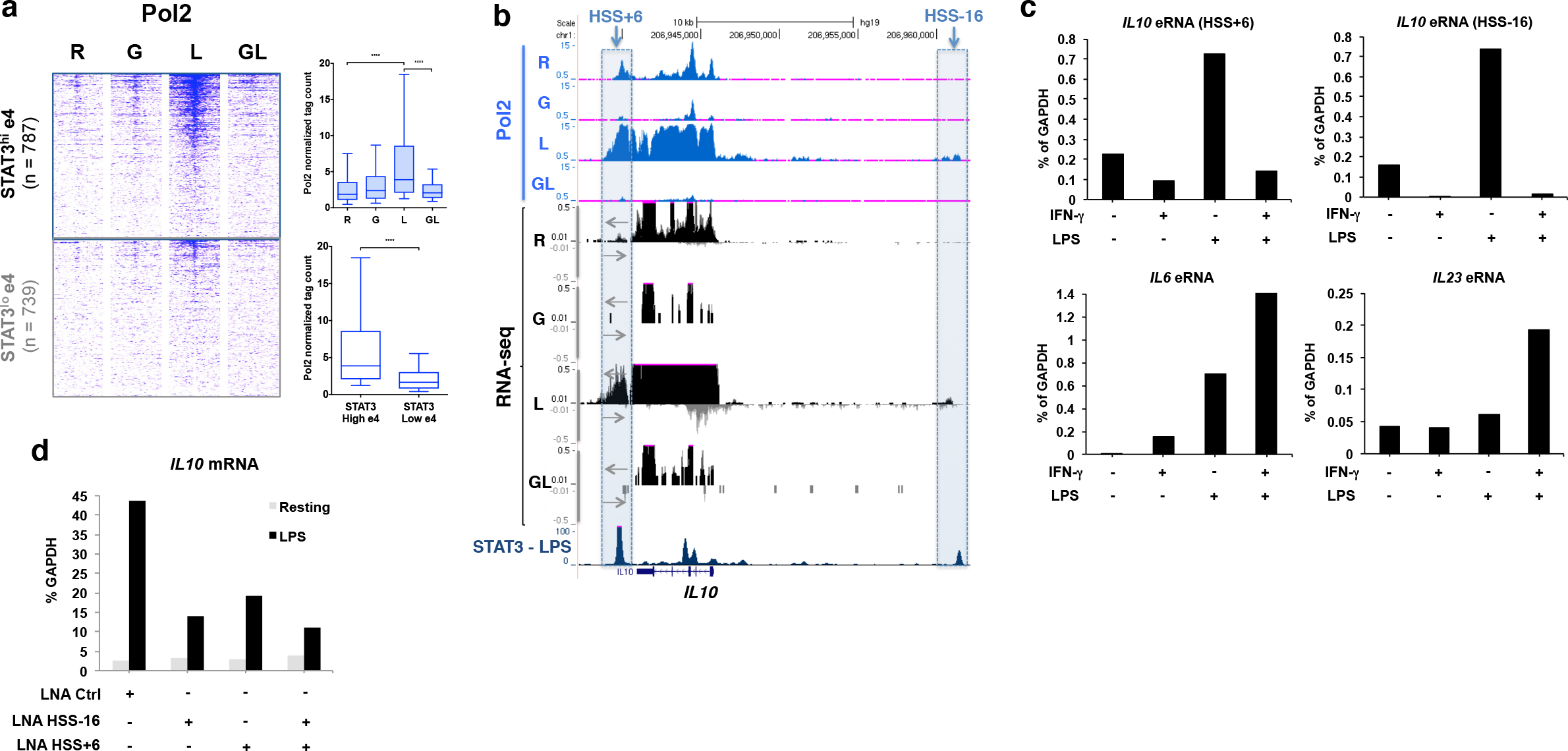
IFN-γ-mediated functional deactivation of STAT3-bound e4 enhancers. (**a**) Heatmaps of Pol II ChIP-seq signals at STAT3^hi^e4 and STAT3^lo^e4 enhancers (defined in **Fig. 6a**). The boxplots indicate normalized tag counts at STAT3^hi^e4 enhancers in the four indicated conditions (top) and at STAT3^hi^e4 and STAT3^lo^e4 enhancers in LPS-stimulated macrophages (bottom). ****p < 0.0001, paired-samples Wilcoxon signed-rank test. (**b**) Representative Genome Browser tracks showing RNA polymerase II (Pol II) occupancy, strand-specific RNA transcripts, and STAT3 occupancy (LPS condition) at enhancers of *IL10*. Boxes enclose HSS+6 (left) and HSS−16 (right). (**c**) RT-qPCR analysis of enhancer RNA (eRNA) expression at two e4 enhancers (top, *IL10*-HSS+6 and *IL-10*-HSS-16) and two e5 enhancers (bottom, *IL6* and *IL23A*). Data are representative of three independent experiments. (**d**) RT–qPCR analysis of *IL10* mRNA in resting and LPS-stimulated macrophages transfected with the indicated LNAs (*IL10*-HSS−16 eRNA and and *IL10*-HSS+6 eRNA). Data are representative of two independent experiments.

## Discussion

Investigation of IFN-γ transcriptional responses has primarily focused on gene activation during ‘M1-like’ macrophage polarization, which is linked to the ‘IFN-γ signature’ observed in autoimmune diseases, and on synergistic activation of inflammatory genes in cooperation with inflammatory factors such as TLR ligands^7,10,42,43^. However, negative regulation of TLR-induced gene expression, and its potential functional consequences and underlying mechanisms are not well understood. This study used a combined transcriptomic and epigenomic approaches to obtain several new insights into negative regulation of TLR-induced gene expression by IFN-γ. First, IFN-γ selectively represses induction of IL-10-inducible anti-inflammatory genes and also metabolic genes that are regulated by a distinct mechanism. Suppression of IL-10-mediated feedback serves to amplify inflammatory gene activation, whereas suppression of metabolic genes, for example those involved in fatty acid metabolism that is linked with M2 macrophages, may promote M1-like polarization. Second, IFN-γ inhibits TLR4-induced genes by targeting associated TLR4-activated enhancers to suppress histone acetylation. Third, IFN-γ inhibits at least two distinct subsets of enhancers, one of which is characterized by enrichment of STAT binding motifs and TLR-induced recruitment of STAT3, CDK8 and cohesin. Another suppressed enhancer subset is enriched for AP-1 motifs and may regulate metabolic genes. Fourth, suppressed enhancers differ from enhancers associated with superactivated ‘synergy’ genes as the latter harbor IRF instead of STAT motifs and bind STAT1 instead of STAT3. However, TLR-inducible enhancers that are modulated by IFN-γ share the property of CDK8 occupancy, suggestive of binding by a select CDK8-containing Mediator complex. Our findings provide insights into previously unexplored mechanisms that selectively regulate TLR responses to promote inflammatory gene activation and M1-like macrophage polarization.

IFN-γ represses basal gene expression in resting macrophages by two distinct epigenetic mechanisms: (1) induction of the negative histone mark H3K27me3 by recruiting EZH2 to promoters^24^ and (2) suppression of the active enhancer histone mark H3K27ac by inhibiting key enhancer-occupying transcription factors, such as MAF^25^. Only a small number of genes is regulated by IFN-γ-mediated direct deposition of H3K27me3, and we did not detect this inhibitory mechanism in the suppression of LPS-induced gene expression. Other groups have suggested that the transcription factors induced by different stimuli, such as IL-4-STAT6^27^ and nuclear receptors (PPARs and LXRs) can directly suppress gene transcription^44^. However, our data did not support a direct repressive role for IFN-γ-induced STAT1 binding at LPS-activated *cis*-regulatory elements, such as promoters and enhancers. Although we cannot rule out the possibility of STAT1 functioning as a transcriptional repressor, it is more likely that IFN-γ mainly mediates the suppression of gene expression by more indirect mechanisms such as STAT1-induced expression of inhibitors of signaling or transcription. Future work is needed to address whether IFN-γ-induced STAT1 has direct repressive functions in other systems.

In macrophages, it has been shown that IL-10, *via* STAT3, plays a pivotal role in the induction of anti-inflammatory factors, such as *BCL3*, *IL4R*, and *TNIP3*^45^, and in regulating an mTORC1-mediated metabolic program including DDIT4^46^. Thus, inactivation of the LPS-induced IL-10-STAT3 negative regulatory pathway by IFN-γ can contribute to ‘uncontrolled’ chronic inflammatory responses. Our epigenomic analysis has identified previously uncharacterized LPS-activated enhancer regions (cluster e4), which are selectively inhibited by IFN-γ. Notably, IFN-γ suppressed LPS-inducible STAT3 binding at e4 enhancers that showed loss of active enhancer marks including histone acetylation and of occupancy by CDK8-cohesin and eRNA-related RNA polymerase II. The e4 enhancers with strong recruitment of STAT3 (STAT3^hi^ e4) showed substantial overlap with enhancers that bind STAT3 as part of the canonical IL-10-induced anti-inflammatory program; a similar overlap was observed between e4-associated and IL-10-inducible genes. Given that LPS-inducible STAT3 binding is related to IL-10-induced anti-inflammatory programs, our findings may provide additional insight into the paradoxical effect of Jak inhibitors on the LPS response, namely that they inhibit STAT3-mediated feedback inhibition^47^. A distinct subset of e4 enhancers with low or minimal STAT3 binding after LPS stimulation (STAT3^lo^ e4) were associated with a distinct IL-10-independent gene set with different functions, such as MAPK signaling pathway and lipid metabolism. We gained new insights into how IFN-γ negatively regulates the expression of LPS-induced autocrine IL-10, which is a major direct inducer of STAT3 activation in response to LPS. Previous work has shown that IFN-γ inhibits TLR-induced autocrine IL-10 production by suppressing AP-1 related pathway, which is related to the differential regulation of MAPK and GSK3 activity^48^. The current study extends this work to suggest that negative regulation of signaling leads to suppression of LPS-induced *IL10* expression *via* epigenetic deactivation of enhancers at the *IL10* locus.

In addition to inhibiting the LPS-induced IL-10-STAT3 autocrine feedback loop, IFN-γ suppressed induction of other multiple additional components of the LPS response that are independent of IL-10. In line with previous work^9^, IFN-γ inhibited transcription of genes that are involved in the ribosomal RNA processing and protein stabilization, and downstream of mTORC1 signaling and Myc. A recent study utilizing co-stimulation of mouse macrophages with IFN-γ and IL-4 suggested that Myc is associated with a component of the IL-4 response that is resistant to suppression by IFN-γ^26^. Our findings suggest that prolonged exposure to IFN-γ that suppresses Myc may overcome this resistance mechanism. mTORC1 and Myc are major regulators of cellular metabolism, and IFN-γ repressed genes involved in various metabolic processes, such as iron transport, purine synthesis, tryptophan metabolism, and lipid metabolism. Of these, lipid metabolism has been linked with M2-like polarization, tryptophan metabolites in anti-inflammatory responses, and iron transport in host defense. Future work will be needed to determine how regulation of these IL-10-independent TLR4-induced metabolic pathways by IFN-γ contributes to the M1 polarized phenotype^49–51^. Overall, IFN-γ-mediated inhibition of anti-inflammatory, translational, and metabolic components of the LPS response is likely coordinated to enable a fully ‘classically activated’ macrophage phenotype.

Our findings highlight differences in function and mode of regulatory element binding between STAT1 and STAT3 during macrophage polarization and LPS challenge. Previous work has shown that during IFN-γ-mediated priming/polarization of macrophages, the genomic profile of STAT1 binding changes from STAT sites to IRF sites, which STAT1 can occupy as part of complexes with IRF1 or (after LPS stimulation) as part of the ISGF3 complex^8,20^. In line with these reports, we found that STAT1 binds to IRF sites coordinately with IRF1 in e5 enhancers, and that joint stimulation with IFN-γ + LPS increased STAT1 binding, which correlated with increased enhancer activity (as assessed by histone acetylation) and superinduction of associated cluster V genes. In contrast, STAT3 bound to e4 enhancers that are enriched for STAT motifs, and joint stimulation with IFN-γ + LPS decreased STAT3 binding, which correlated with decreased enhancer activity and suppression of associated cluster IV genes. These results support a model whereby intrinsic DNA sequences (TF binding motifs) in genomic regulatory elements determine whether IFN-γ primes or suppresses activation of an enhancer by LPS. Namely, in IFN-γ-primed macrophages enhancers that contain IRF but not STAT sites (e.g. e5 enhancers) are subject to regulation mostly by STAT1 and exhibit enhanced LPS responses, whereas enhancers that also contain a STAT site are regulated by STAT3 and exhibit attenuated LPS responses. Motif analysis also found that AP-1 motif was highly enriched at STAT3-bound enhancers, which are suppressed by IFN-γ, suggesting that IFN-γ-regulated AP-1 transcription factors including BATF and FOSL2 might serve as ‘auxiliary’ transcription factors and cooperate with STAT3 to regulate e4 enhancers. Identifying additional auxiliary transcription factors that cooperate with STAT1 and STAT3 to drive expression of different LPS-regulated gene sets may provide further insight into STATs-dependent transcriptional regulation in a gene-specific manner.

In summary, this study provides insights about how two stimuli can cooperate at the epigenomic level to achieve differential regulation of gene sets with distinct and even opposing functions. In the case of IFN-γ and LPS, differential regulation of enhancers with different sequence architecture results in decreased expression of anti-inflammatory and metabolic genes, with superinduction of inflammatory genes that may enhance inflammatory responses. The distinct enhancer classes can potentially be targeted to restrain excessive inflammation.

## Methods

### Cell culture

Peripheral blood mononuclear cells were obtained from blood leukocyte preparations purchased from the New York Blood Center by density gradient centrifugation with Ficoll (Thermo Fisher Scientific) using a protocol approved by the Hospital for Special Surgery Institutional Review Board. Primary human CD14^+^ monocytes were obtained from peripheral blood, using anti-CD14 magnetic beads, as recommended by the manufacturer (Miltenyi Biotec). Monocytes were cultured in RPMI 1640 medium (Invitrogen) supplemented with 10% heat-inactivated defined FBS (HyClone Fisher), penicillin/streptomycin (Invitrogen), L-glutamine (Invitrogen), and 10 ng/ml human macrophage colony-stimulating factor (M-CSF; Peprotech) in the presence or absence of 100 U/ml human IFN-γ (Roche) as indicated; IFN-γ was added at the same time as M-CSF at initiation of cultures. LPS was purchased from Invivogen (tlrl-3pelps).

### Analysis of RNA

Total RNA was extracted from cells using RNeasy Mini kit (QIAGEN), and 500 ng of total RNA was reverse transcribed using the RevertAid First Strand cDNA Synthesis kit (Fermentas). Real-time PCR was performed in triplicate with Fast SYBR Green Master Mix and 7500 Fast Real-time PCR system (Applied Biosystems). Primer sequences are provided in the **Supplementary Table 1**.

### RNA-seq

After RNA extraction, libraries for sequencing were prepared using the Illumina TruSeq RNA Library Prep Kit following the manufacturer’s instructions. High throughput sequencing (50 bp, paired-end) was performed at the Genomics Resources Core Facility of Weill Cornell Medicine. On average 100 million reads were obtained per sample. Sequenced reads were mapped to reference human genome (hg19 assembly) using STAR aligner^52^ with default parameters and Cufflinks version 2.2.1^53^ was used to estimate the abundance of transcripts. The expression levels of genes in each sample were normalized by means of fragments per kilobase of transcript per million mapped reads (FPKM). The concordance between replicates was very high (R^2^ range, 0.943-0.964).

### RNA-seq analysis

Differentially expressed genes (DEGs) were identified using edgeR v3.16.5 ^54,55^. Read counts for edgeR analysis were obtained with featureCounts v1.5.1 ^56^. After eliminating absent features (zero counts), the raw counts were normalized with edgeR, followed by differential expression analysis. Significantly up-or down-regulated genes were defined as expressed genes with *p*-value adjusted for multiple testing (FDR < 0.01) and log_2_ fold-change of at least 1. To generate the heatmap of K-mean clusters, we used GENE-E (Broad Institute) set to global comparison and average-centered. The value of K was chosen at 6 because lower values failed to identify all meaningful clusters and higher values subdivided meaningful clusters. To find the GO terms enriched in differentially regulated genes, we used the DAVID web-tool^57^ and Gene Set Enrichment Analysis (GSEA) of MSigDB gene sets (allmark, KEGG, and REACTOME).

### Analysis of gene expression in IL-10 stimulated human monocytes

Microarray data sets were retrieved from GSE43700. The raw data were normalized by a quantile normalization method using the preprocessCore package in R. Normalized expression levels were averaged within the same condition and fold-change of the average for each gene was calculated.

### ChIP and ChIP-seq

Cells were crosslinked for 5 min at room temperature by the addition of one-tenth of the volume of 11% formaldehyde solution (11% formaldehyde, 50 mM HEPES pH 7.5, 100 mM NaCl, 1 mM EDTA pH 8.0, 0.5 mM EGTA pH 8.0) to the growth media followed by 5 min quenching with 100 mM glycine. Cells were pelleted at 4°C and washed with ice-cold PBS. The crosslinked cells were lysed with lysis buffer (50 mM HEPES-KOH pH 7.5, 140 mM NaCl, 1 mM EDTA, 10% glycerol, 0.5% NP-40, and 0.25% Triton X-100) with protease inhibitors on ice for 10 min and washed with washing buffer (10 mM Tris-HCl, pH 8.0, 200 mM NaCl, 1 mM EDTA, 0.5 mM EGTA) for 10 min. The lysis samples were resuspended and sonicated in sonication buffer (10 mM Tris-HCl, pH 8.0, 100 mM NaCl, 1 mM EDTA, 0.5 mM EGTA, 0.1% Na-Deoxycholate, 0.5% N-lauroylsarcosine) using a Bioruptor (Diagenode) with 30 sec ON, 30 sec OFF on high power output for 18 cycles. After sonication, samples were centrifuged at 14,000 rpm for 10 minutes at 4°C and 5% of sonicated cell extracts were saved as input. The resulting whole-cell extract was incubated with Protein A Agarose for ChIP (EMD Millipore) for 1 hr at 4°C. Precleared extracts were then incubated with 50 μl (50% v/v) of Protein A Agarose for ChIP (EMD Millipore) with 5 μg of the appropriate antibody overnight at 4°C. ChIP lysates were generated from 2 x 10^7^ to 3 x 10^7^ cells (for PU.1, C/EBPβ, STAT3, c-Jun, and RNA Polymerase II ChIP) or 10 x 10^7^ cells (for p300, MED1, CDK8, and β SMC1 ChIP) respectively. ChIP antibodies against PU.1 (sc-352), C/EBPβ (sc-150), STAT3 (sc-482), c-Jun (sc-1694), p300 (sc-585) and CDK8 (sc-1521) were from Santa Cruz Biotechnology. Antibodies against RNA Polymerase II (MMS-126R) were from Covance. Antibodies against MED1 (A300-793A) and SMC1 (A300-055A) were from Bethyl Laboratories. After overnight incubation, beads were washed twice with sonication buffer, once with sonication buffer with 500 mM NaCl, once with LiCl wash buffer (10 mM Tris-HCl pH 8.0, 1 mM EDTA, 250 mM LiCl, 1% NP-40), and once with TE with 50 mM NaCl. DNA was eluted in freshly prepared elution buffer (1% SDS, 0.1M NaHCO3). Cross-links were reversed by overnight incubation at 65°C. RNA and protein were digested using RNase A and Proteinase K, respectively and DNA was purified with ChIP DNA Clean & Concentrator^™^ (Zymo Research). For ChIP-qPCR assays, immunoprecipitated DNA was analyzed by quantitative real-time PCR and normalized relative to input DNA amount.

For ChIP-seq experiments, 10 ng of purified ChIP DNA per sample were ligated with adaptors and 100-300 bp DNA fragments were purified to prepare DNA libraries using Illumina TruSeq ChIP Library Prep Kit following the manufacturer’s instructions. ChIP libraries were sequenced (50 bp single end reads) using an Illumina HiSeq 2500 Sequencer at the Epigenomic Core Facility of Weill Cornell Medicine per manufacturer’s recommended protocol. Because of limitations on cell numbers and to decrease variability related to differences among individual donors, chromatin immunoprecipitations were performed using pooled samples from more than two (for STAT3) or four (for CDK8 and SMC1) different donors. For STAT3, a second experiment with pooled samples from several donors was performed and congruence between the replicates was assessed by generating scatter plots and estimating Pearson correlation coefficients (**Figure S4A**). After ascertaining close correlation between replicates, we performed bioinformatic analysis using replicate 1 and confirmed key results using replicate 2. The H3K27ac, STAT1 and IRF1 data were from GSE43036.

### ATAC-seq

ATAC-seq was performed as previously described^58^. 50,000 cells were centrifuged at 500 g for 5 min at 4°C. Cell pellets were washed once with 1x PBS and cells were pelleted by centrifugation using the previous settings. Cell pellets were resuspended in 25 μl of lysis buffer (10 mM Tris-HCl pH 7.4, 10 mM NaCl, 3 mM MgCl_2_, 0.1% IGEPAL CA-630) and centrifuged immediately 500 g for 10 min at 4°C. The cell pellet was resuspended in the transposase reaction mix (25 μL 2× TD buffer (Nextera DNA Sample Preparation Kit), 2.5 μL Illumina Tn5 transposase and 22.5 μL nuclease-free water). The transposition reaction was carried out for 30 min at 37°C. Directly following transposition, the sample was purified using a QIAGEN MinElute Purification Kit. Then, we amplified library fragments using NEBNext 2x PCR master mix and 1.25 M of Nextera PCR primers, using the following PCR conditions: 72 °C for 5 min; 98 °C for 30 s; and thermocycling at 98°C for 10 s, 63°C for 30 s and 72°C for 1 min. The libraries were purified using a QIAGEN PCR purification kit yielding a final library concentration of ~30 nM in 20 μL. Libraries were amplified for a total of 10-13 cycles and were subjected to high-throughput sequencing using the Illumina HiSeq 2500 Sequencer.

### ChIP-seq and ATAC-seq analysis

For ChIP-seq and ATAC-seq experiments, sequenced reads were aligned to reference human genome (GRCh37/hg19 assembly) using Bowtie2 version 2.2.6 (Langmead and Salzberg, 2012) with default parameters, and clonal reads were removed from further analysis. A minimum of 20 million uniquely mapped reads were obtained for each condition. We used the makeTagDirectory followed by findPeaks command from HOMER version 4.9.1 (Heinz et al.,2010) to identify peaks of ChIP-seq enrichment over background. A false discovery rate (FDR) threshold of 0.001 was used for all data sets. Regions that overlap with blacklist identified by the ENCODE project were filtered out. The total number of mapped reads in each sample was normalized to ten million mapped reads. ChIP-seq data were visualized by preparing custom tracks for the UCSC Genome browser. For identifying enhancer regions that were differentially acetylated in at least one of four conditions (FDR < 0.05, >2-fold changes in any pairwise comparison), we used the getDifferentialPeaksReplicates.pl with parameters - genome hg19 -balanced -t H3K27ac_L1/ H3K27ac_L2/ -b H3K27ac_R1/ H3K27ac_R2/ -p enhancers.bed from HOMER package and then merged using mergePeaks - size given. For the clustering of enhancers, we used GENE-E (Broad Institute) set to global comparison and average-centered. The value of K was chosen at 5 because lower values failed to identify all meaningful clusters and higher values subdivided existing clusters. The third enhancer cluster (C3) in **Supplementary Fig. 2c** was further classified into two clusters (e2 and e3) in **Fig.2a** based on the changes in H3K27ac between IFN-γ-stimulated (G) and IFN-γ-primed LPS-stimulated (GL) macrophages. For the distribution plot of ChIP-seq signals in **Fig.4c** and g, we used the annotatePeaks.pl command with parameters-size 2000-hist 10 to generate histograms for the average distribution of normalized tag densities. For functional annotations of the enhancers, enriched GO Biological Process and MSigDB Pathways were compiled from the GREAT version 3.0.0 (McLean et al., 2010) on each subset of enhancer-associated genes. GO and MSigDB pathways were ranked based on p-values.

### Motif enrichment analysis

*de novo* and known motif analysis was performed on ±100 bp centered on the ATAC-seq peak that overlapped either PU.1 or C/EBPβ using command “findMotifsGenome.pl” from HOMER package. Peak sequences were compared to random genomic fragments of the same size and normalized G+C content to identify motifs enriched in the targeted sequences.

### LNA transfection

LNAs (LNA^™^ longRNA GapmeR) were designed and synthesized by Exiqon. Knockdown experiments with LNAs were performed using Human Monocyte Nucleofector buffer (Lonza Cologne) and the AMAXA Nucleofector System program Y001 for human monocytes according to the manufacturer’s instructions. Two different LNA^™^ longRNA GapmeRs (negative control A and B) were used as control. Cells were harvested for RNA analysis 24 h after transfection.

The following LNAs were used for knockdown studies of *IL10* eRNAs. Custom LNA for *IL10* eRNA (HSS-16): ATAGAGAGGAGATGCA, GCAGTCTAGCTTGGTG. Custom LNA for *IL10* eRNA (HSS+6): GGATTTGGCGGGAGTT, TCCTAGTGCCAGAAGC.

### Statistical analysis

Statistical tests were selected based on appropriate assumptions with respect to data distribution and variance characteristics. Wilcoxon signed-rank test or two-tailed paired t-test was used for the statistical analysis of two paired samples. Welch’s t-test or unpaired Student’s t-test was used for the statistical analysis of two non-paired samples. Statistical significance was defined as p < 0.05. The whiskers of box plots represent the 10–90th percentiles of the data. Statistical analyses were performed using GraphPad Prism 7.

### Data availability statement

RNA-seq, ChIP-seq and ATAC-seq data for this project have been deposited at NCBI’s Gene Expression Omnibus (GEO) and GSE number is pending.

---

**Supplementary Fig. 1.**
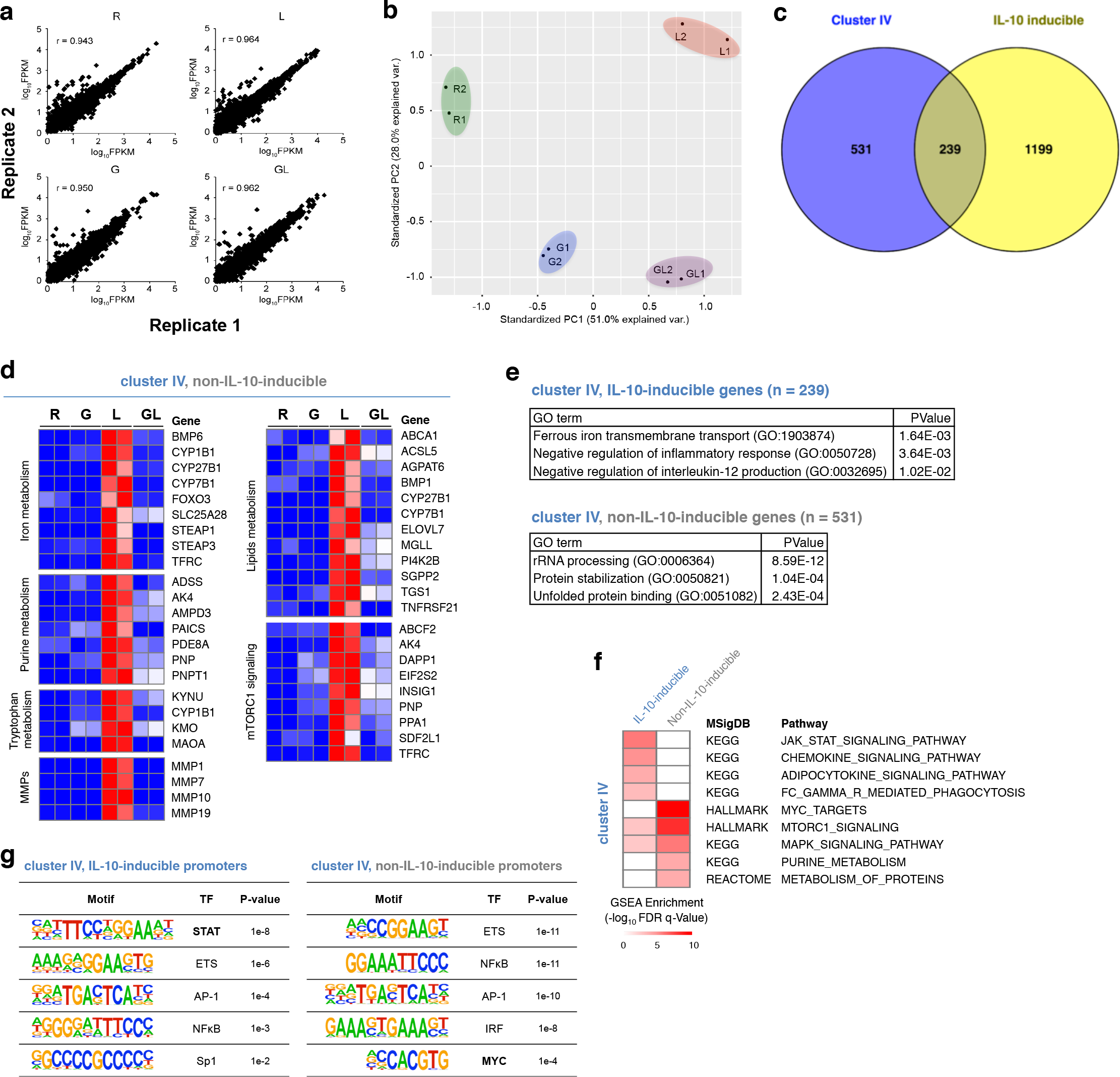
Transcriptomic profiling of IFN-γ-primed LPS-stimulated macrophages. (**a**) Correlation of RNA-seq replicates. (**b**) Principle Component Analysis (PCA) for RNA-seq replicates among four conditions. (**c**) Venn diagram showing number of genes in cluster IV (n = 531, blue), genes induced by IL-10 alone (n = 1199, yellow), and the overlap between cluster IV and IL-10-inducible genes (n = 239). (**d**) Heatmap showing the relative expression of examples of cluster IV genes that are not inducible by IL-10. Distinct biological functions showed on the left. (**e**) Gene ontology (GO) analysis using DAVID 6.8. (**f**) Heat map showing the FDR q-value significance of GSEA enrichment for the two classes of cluster IV genes classified in (**c**). (**g**) Known motif analysis of promoters. Data is representative of two experiments.

**Supplementary Fig. 2.**
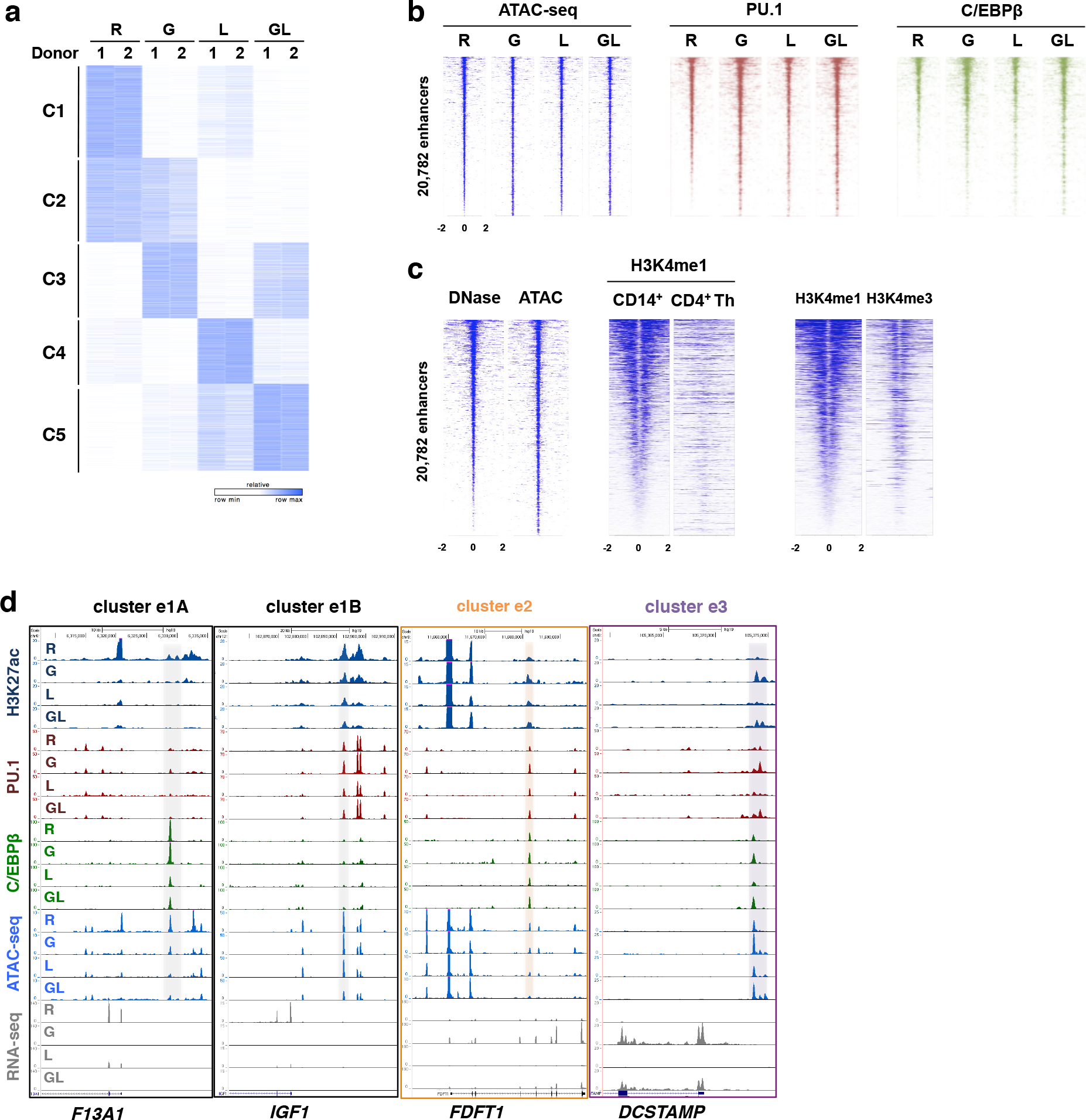
IFN-γ regulates TLR4-activated enhancer landscapes. (**a**) K-means clustering (K = 5) of H3K27ac ChIP-seq intensity in open chromatin regions (ATAC-seq peaks) that were bound by PU.1 or C/EBPβ. (**b**) Heatmaps of ATAC-seq, PU.1, and C/EBPβ ChIP-seq signals at 20,782 enhancers differentially regulated at the H3K27-Ac level. (**c**) Left panels show congruence of ATAC-seq peaks with DNase-seq peaks from ENCODE project. Heatmaps of normalized tag densities for DNase-seq (CD14^+^ monocytes from ENCODE, GSM1024791) and ATAC-seq (monocyte-derived macrophages used in this project) at the 20,782 enhancers defined and analyzed in this project. Middle panels show heatmaps of normalized tag densities for H3K4me1 ChIP-seq from CD14+ monocytes (left, GSM1102793) and CD4^+^ T cells (right, GSM1220567) at the 20,782 enhancers. Right panels show heatmaps of normalized tag densities for H3K4me1 ChIP-seq from CD14+ monocytes (left) and H3K4me3 ChIP-seq from CD14+ monocytes (right, GSM945225) at the 20,782 enhancers. Four-kilobase windows are shown centered at the midpoints of the ATAC-seq peak. (**d**) Representative UCSC Genome Browser tracks displaying normalized tag-density profiles at enhancers of *F13A1, IGF1, FDFT1,* and *DCSTAMP* in the four indicated conditions. Boxes enclose cluster e1A, e1B, e2, and e3 enhancers.

**Supplementary Fig. 3.**
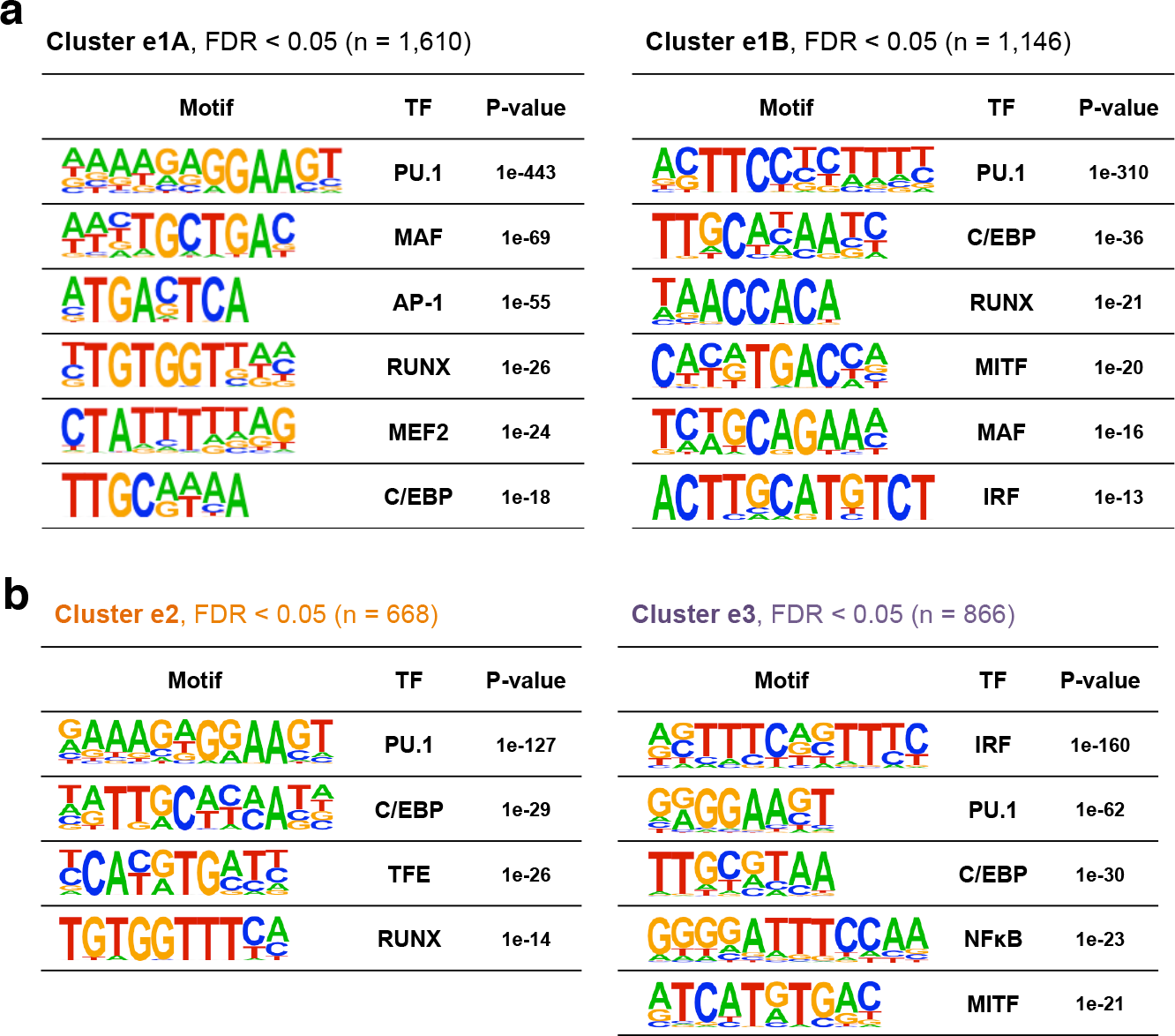
Distinct motif enrichment in different enhancer clusters. (**a**) The most significantly enriched transcription factor (TF) motifs identified by *de novo* motif analysis using HOMER in cluster e1A (left) and cluster e1B (right) enhancers. (**b**) The most significantly enriched transcription factor (TF) motifs identified by *de novo* motif analysis using HOMER at cluster e2 (left) and cluster e3 (right) enhancers.

**Supplementary Fig. 4.**
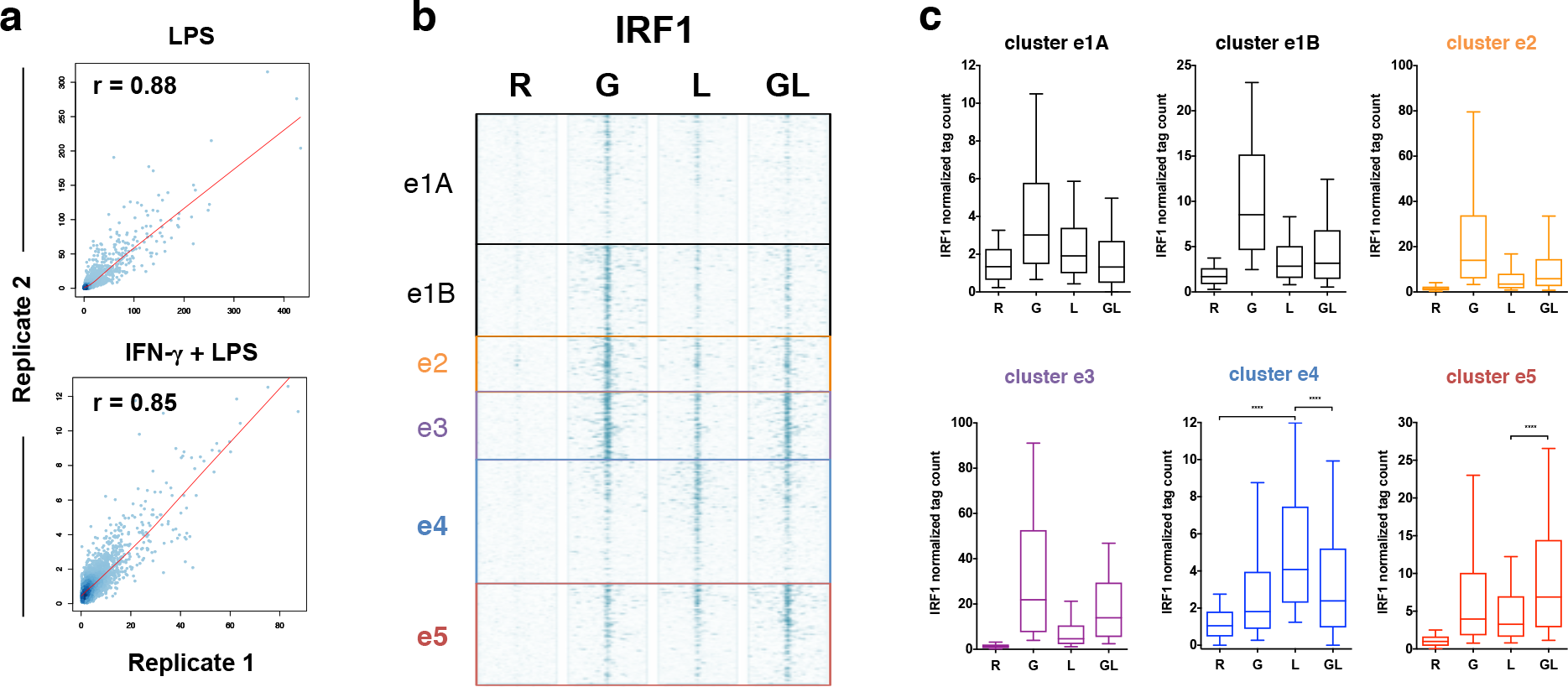
Asymmetric TF binding at each enhancer cluster. (**a**) Correlation of STAT3 ChIP-seq replicates in LPS-stimulated (top) and IFN-γ-primed LPS-stimulated macrophages (bottom). (**b** Heatmaps showing IRF1 ChIP-seq signals at each enhancer cluster defined in **Fig. 2a**. (**c**) The boxplots indicate IRF1 normalized tag counts at each enhancer cluster. ****p < 0.0001, paired-samples Wilcoxon signed-rank test.

**Supplementary Fig. 5.**
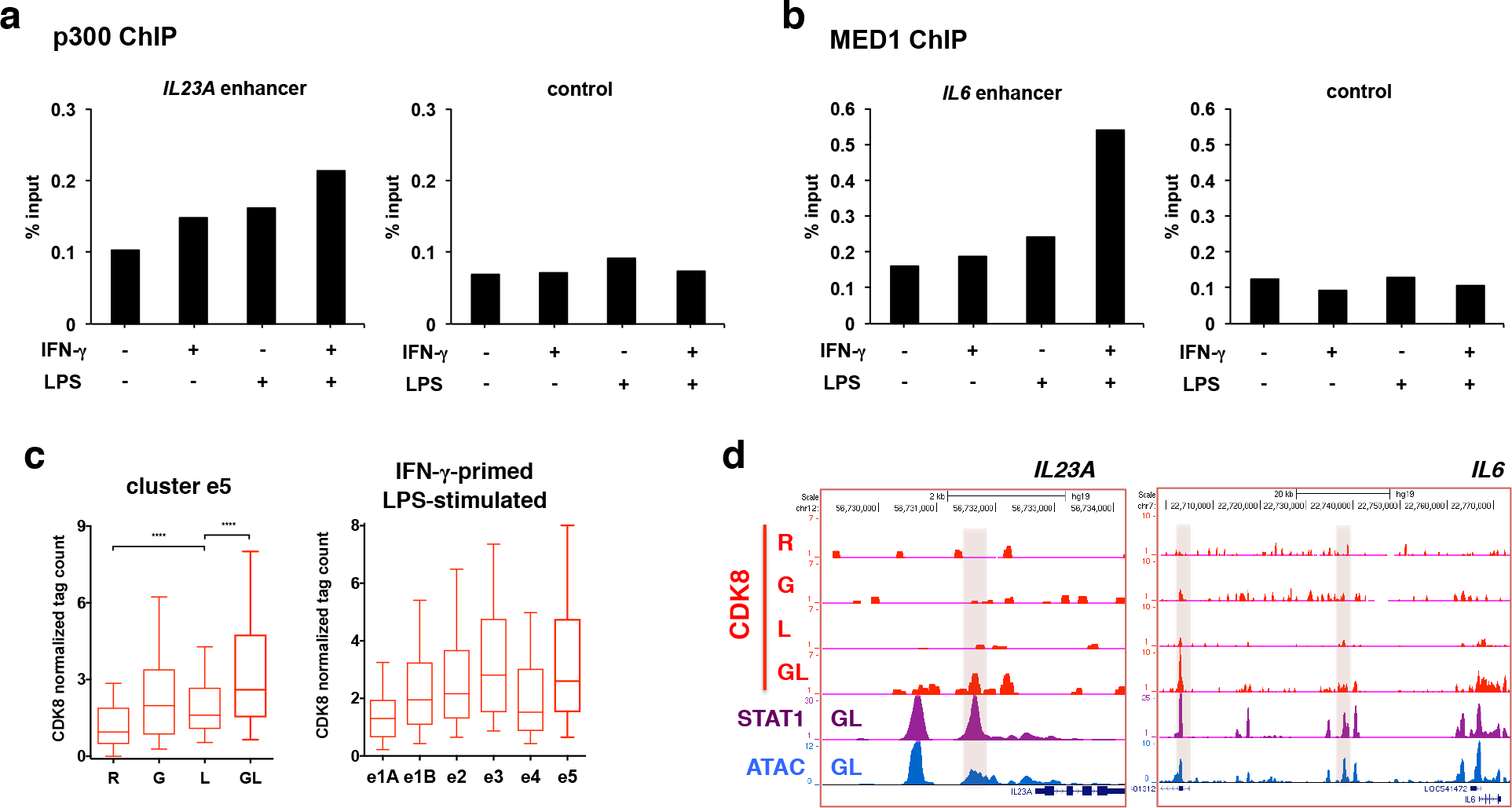
IFN-γ priming increases LPS-induced binding of p300, MED1 and CDK8 at e5 enhancers. (**a**) ChIP-qPCR analysis of p300 occupancy at the e5 enhancer of *IL23A*. (**b**) ChIP-qPCR of MED1 at the e5 enhancer of *IL6*. Data are representative of three independent experiments. (**c**) The boxplot (left) indicates CDK8 normalized tag counts at e5 enhancers in the four indicated conditions. ****p < 0.0001, paired-samples Wilcoxon signed-rank test. The boxplot (right) indicates CDK8 normalized tag counts at each enhancer cluster in IFN-γ-primed LPS-stimulated macrophages. (**d**) Representative UCSC Genome Browser tracks displaying normalized tag-density profiles at e5 enhancers of *IL23A* and *IL6* in the four indicated conditions (CDK8) and the LPS-stimulated condition (STAT1 ChIP-seq and ATAC-seq).

**Supplementary Fig. 6.**
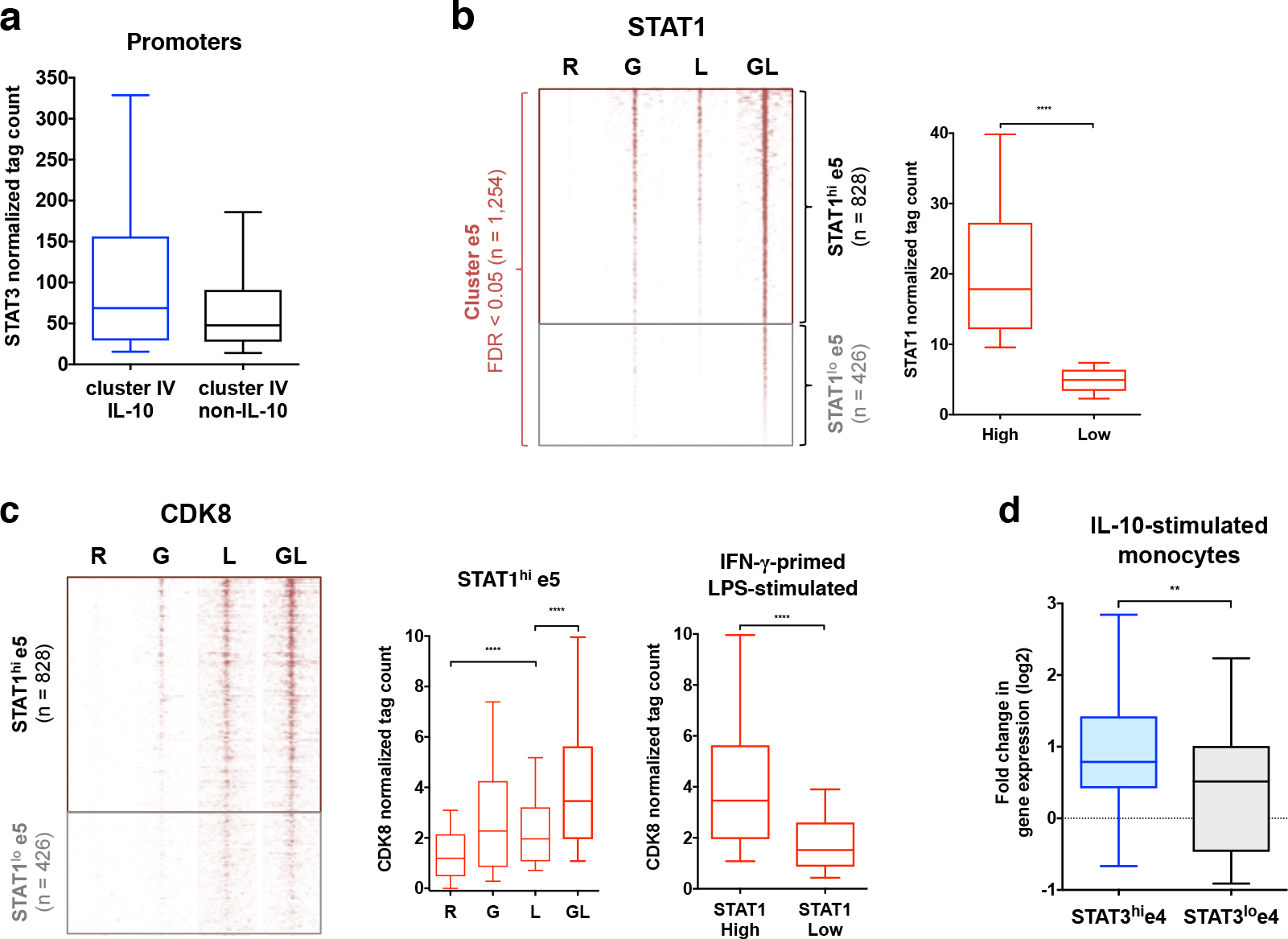
e5 enhancers subdivide into two groups based on coordinate STAT1 and CDK8 occupancy. (**a**) The boxplot indicates normalized tag counts at promoters of IL-10-inducible cluster IV genes (blue) and non-IL-10-inducible cluster IV genes (black). (**b**) Heatmap of STAT1 ChIP-seq signals at cluster e5 enhancers in the four indicated conditions. Enhancers were separated into two subsets: STAT1^hi^e5 (n = 828) and STAT1^lo^e5 (n = 426) based upon a log_2_ normalized tag count cutoff od 3). The boxplot (right) indicates normalized tag counts at STAT1^hi^e5 and STAT1^lo^e5 enhancers. (**c**) Heatmaps of CDK8 ChIP-seq signals at the two subsets of e5 enhancers (defined in **b**). The boxplots indicate normalized tag counts at STAT1^hi^e5 enhancers (left) and at STAT1^hi^e5 and STAT1^lo^e5 enhancers in IFN-γ-primed LPS-stimulated macrophages (right). *p < 0.0001, paired-samples Wilcoxon signed-rank test. (**d**) Boxplots of the change in gene expression after IL-10 stimulation of macrophages of the differentially expressed genes nearest (within 100 kb) to STAT3^hi^e4 (blue) or STAT3^lo^e4 (black) enhancers. **p = 0.0032 by Welch’s t test.

**Supplementary Fig. 7.**
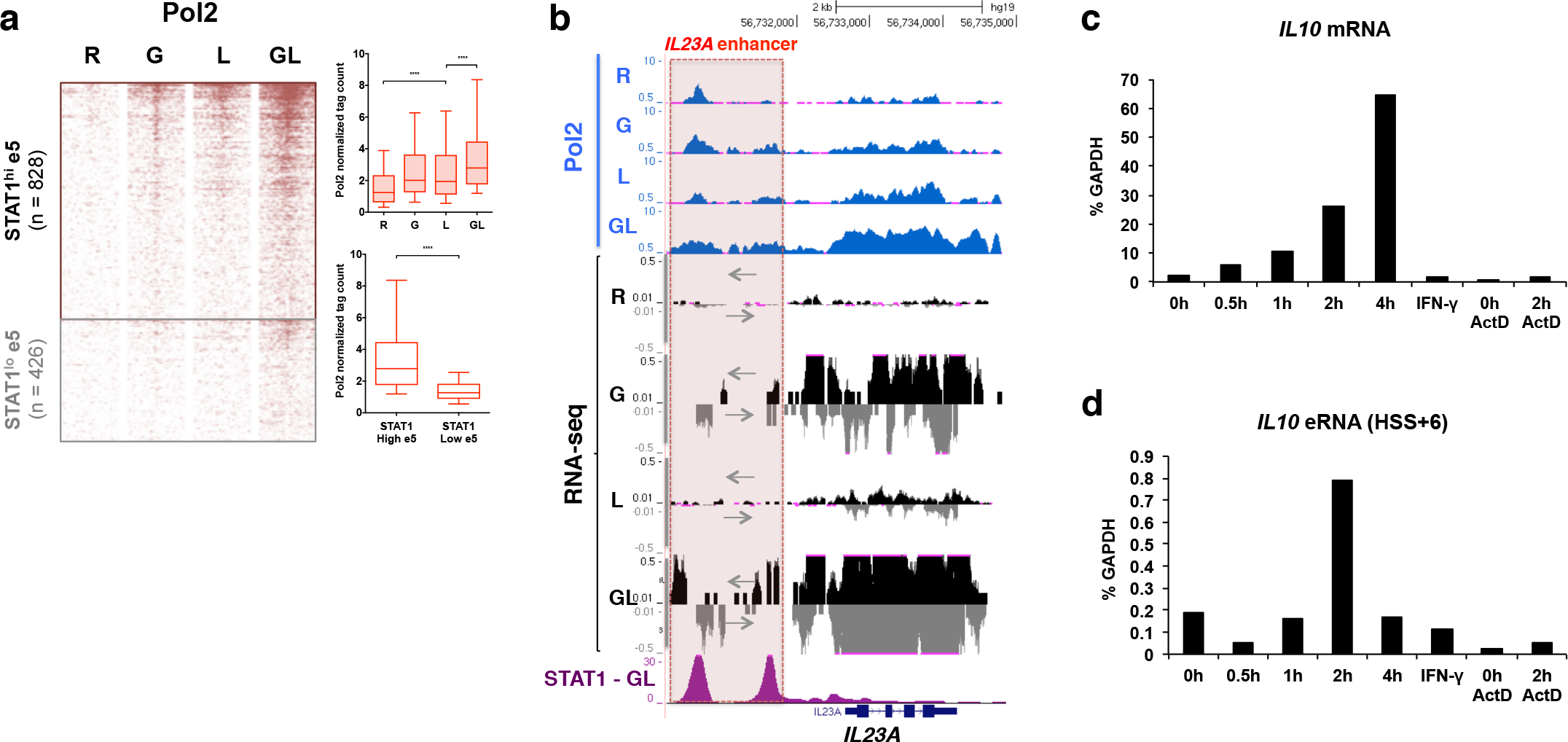
IFN-γ-mediated functional activation of STAT1-bound e5 enhancers. (**a**) Heatmaps of Pol II ChIP-seq signals at STAT1^hi^e5 and STAT1^lo^e5 enhancers (defined in **Supplementary Fig. 6b**) in the four indicated conditions. The boxplots indicate normalized tag counts at STAT1^hi^e5 enhancers (top) and at STAT1^hi^e5 and STAT1^lo^e5 enhancers in IFN-γ-primed LPS-stimulated macrophages (bottom). ****p < 0.0001, paired-samples Wilcoxon signed-rank test. (**b**) Representative Genome Browser tracks showing RNA polymerase II (Pol II) occupancy, strand-specific RNA transcripts, and STAT1 occupancy (IFN-γ + LPS condition) at enhancers of *IL23A*. Box encloses enhancer of *IL23A*. (**c**) RT-qPCR analysis of *IL10* mRNA expression in macrophages cultured without or with IFN-γ for 48 hr and then stimulated with LPS. 5 μg/mL actinomycin D (Act D) was added 1 hr prior to harvesting. (**d**) RT-qPCR analysis of *IL10* eRNA (HSS+6) expression in macrophages cultured without or with IFN-γ for 48 hr and then stimulated with LPS for the indicated time course. 5 μg /mL actinomycin D (ActD) was added 1 hr prior to harvesting. Data are representative of two independent experiments.

**Supplementary Table 1.**
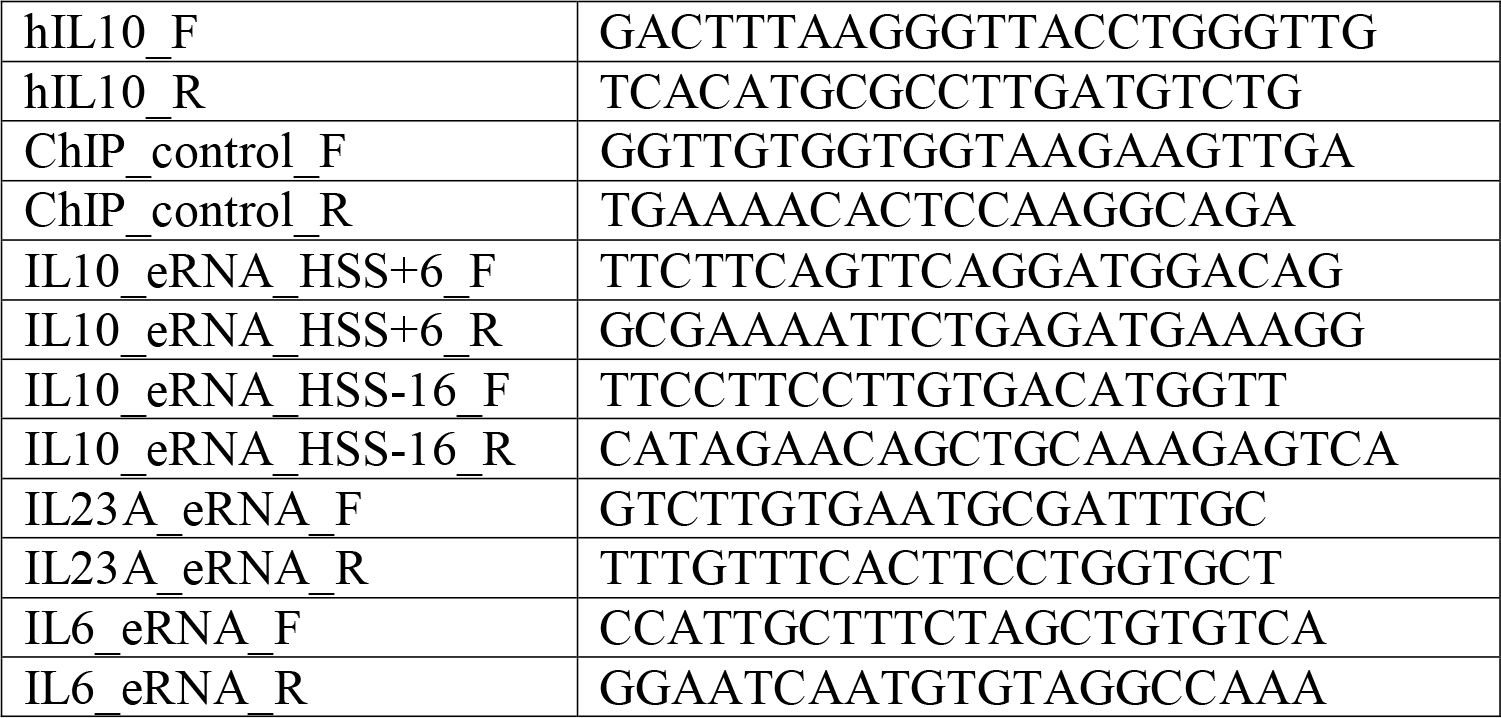
Primers for expression of mRNA or eRNA and ChIP-qPCR.

